# Isoform-specific functions of Numb in breast cancer progression, metastasis and proteome remodeling

**DOI:** 10.1101/2021.02.01.429237

**Authors:** Yangjing Zhang, Sascha E. Dho, Craig D. Simpson, Kamal Othman, Andrew Bondoc, C. Jane McGlade

## Abstract

Deregulated alternative splicing of the endocytic adaptor *NUMB* resulting in high expression of Exon9in (exon 9-containing) isoforms has been reported in several cancer types. However, the role of Numb isoform expression in tumor progression and the underlying mechanisms remain elusive. Here, we report greater exon 9 inclusion in multiple cancer types including all subtypes of breast cancer, and correlation of higher exon 9 inclusion in patients with worse prognosis. Deletion of Exon9in in breast cancer cells leads to reduced cell growth and a significant decrease of lung metastasis in orthotopic xenograft experiments. Quantitative mass spectrometry revealed downregulation of proteins involved in EMT and ECM organization and remodeling of the endocytic protein network in cells lacking the Exon9in Numb isoforms. Exon 9 deletion also results in reduced surface levels of ITGβ5, and downstream signaling to ERK and SRC, consistent with enhance lysosomal targeting mediated by the remaining Exon9sk (exon 9 skipping) Numb isoforms.

**SIGNIFICANCE:** Expression of *NUMB* Exon9in protein isoforms correlate with worse progression free survival, particularly in breast cancer. Our findings also reveal that Exon9in isoforms promote breast cancer progression by relieving Numb mediated down regulation of integrins and implicate Numb alternative splicing as a progression factor in multiple cancer types.

## Introduction

*NUMB* encodes a cell fate determinant (1) and conserved adaptor (2, 3) that binds to endocytic proteins and regulates the trafficking of surface proteins such as Notch (4), Integrins (5, 6) and E-cadherin (7, 8). Numb also promotes proteasomal degradation of Gli1 (9) and the stabilization of p53 (10). Although Numb is not known to be mutated in cancer, loss of protein expression that abrogates these functions has led to its characterization as a tumor suppressor (9, 11, 12). However, this conclusion has been complicated by more recent studies of functionally distinct Numb isoforms. While earlier studies implicated reduced Numb protein levels in breast and lung cancer (11, 12), subsequent studies revealed that changes in Numb alternative splicing is a frequent event, particularly in lung cancer (13).

In vertebrates, four Numb isoforms are expressed based on the alternative splicing of two cassette exons: exon 3 and exon 9 (14). The alternative splicing is developmentally regulated in rat and human brain (15), mouse cerebral cortex (16), retina (17) and pituitary gland (18), where exon 9 is predominantly included in stem cells and skipped in differentiated cells. Overexpression of exon 9 included (Exon9in) Numb isoforms in mouse stem cells expands the progenitor population (15, 16). Conversely, the expression of exon 9 skipped (Exon9sk) isoforms drives differentiation (15, 16) and the specific knockdown of Exon9sk but not Exon9in in chick spinal cord impairs neural differentiation (19).

Increased expression of Numb exon 9 is a feature of multiple cancer types including cervical squamous cell carcinoma (20), non-small cell lung cancer (13), urothelial carcinoma (21) and hepatocellular carcinoma (22). In breast and lung cancer cell lines, activated MEK/ERK signaling promotes exon 9 inclusion (23). It has also been reported that the RBM5/6 and RBM10 splicing factors differentially regulate *NUMB* exon 9 splicing to control lung cancer cell proliferation (24). These studies suggest that oncogenic pathways activate exon 9 inclusion, and expression of tumor promoting Exon9in Numb isoforms.

The mechanisms by which Exon9in isoforms of Numb contribute to tumorigenesis remain poorly understood. In addition to a potential gain-of-function acquired through increased Exon9in expression, evidence also suggests that the Exon9in isoforms may antagonize the tumor-suppressive activities of Exon9sk Numb proteins. For example, overexpression studies have shown differential effects of Exon9in isoforms on Notch signaling (24, 25). In addition, several studies suggest different roles of Exon9in and Exon9sk isoforms in endocytic protein association (26), and trafficking (27).

Here, we show that the inclusion level of *NUMB* exon 9 is higher in multiple cancer types and that greater exon 9 inclusion is associated with worse prognosis, especially in breast cancer. Deletion of exon 9 in a basal like human breast cancer cell line leads to a reduction in lung metastasis in a xenograft model and decreased cell growth in vitro. Quantitative proteomics revealed that exon 9 deletion alters the expression of proteins involved in epithelial mesenchymal transition (EMT), extracellular matrix (ECM) organization and the endocytic network. Furthermore, cells lacking Exon9in Numb isoforms have decreased surface levels of ITGβ5 and downstream signaling. These findings reveal the mechanisms by which alternative splicing and Numb protein isoform expression contribute to breast cancer progression and metastasis.

## Results

### *NUMB* exon 9 usage is increased in multiple cancer types and correlates with poor prognosis

To compare the expression of Numb isoforms in primary cancer tissue versus normal tissue, we analyzed the 21 out of 32 cancer types from The Cancer Genome Atlas (TCGA) (28) which have matched normal samples (Fig. 1B, Supplementary Fig. S1A-C). The Percent Spliced In Index (PSI) represents the percentage of mRNA transcripts that include a specific exon. Numb exon 9 PSI value was significantly higher in 11/21 cancer types analyzed compared to tissue matched normal, including cervix (increased 56% above normal), uterine (44%), lung (squamous cell carcinoma 30%, adenocarcinoma 39%), breast (31%), bladder (28%), pancreas (23%), prostate (21%), liver (22%), bile duct (20%), and stomach (14%). Since the PSI value is independent of the expression level of the gene, it is possible to cross validate exon 9 inclusion in TCGA tumor samples with the Genotype-Tissue Expression (GTEx) database of normal tissues. It also allows us to include additional cancers in TCGA dataset which had no matched normal tissue. Comparison of TCGA tumor to matched normal tissue from the GTEx database confirmed increased Exon9in Numb isoform expression in 10 out of the 11 cancer types identified in the first analysis (Supplementary Fig. S1D). In addition, this analysis showed significantly higher exon 9 inclusion in ovarian serous cystadenocarcinoma (38% increase) and testicular germ cell tumors (16%), two cancer types which had no matched normal tissue in the TCGA dataset (Supplementary Fig. S1D).

**Figure 1.**
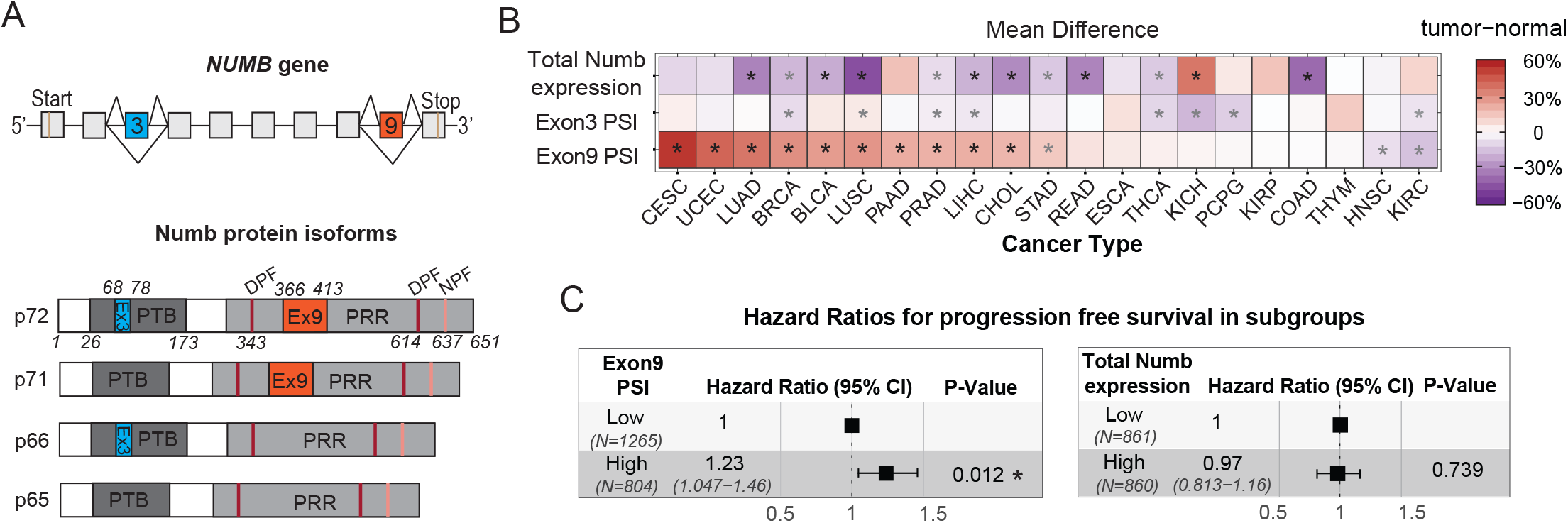
Increased *NUMB* exon 9 inclusion is a common feature in cancer and is associated with worse prognosis. **(A)** Schematic diagrams showing the alternatively spliced exons of the human *NUMB* gene and the domain structure of Numb protein isoforms. The amino acid residues at the boundaries of exon 3 and exon 9 are numbered. Location of phosphotyrosine-binding domain (PTB), proline-rich region (PRR), and tripeptide protein interaction motifs Asn-Pro-Phe (NPF) and Asp-Pro-Phe (DPF) are shown. **(B)** Histogram showing the inclusion levels of *NUMB* exon 9 and exon 3 (represented by the PSI value) and total Numb expression (represented by normalized transcript level in readcounts) in 21 cancer types compared to tissue matched normal samples in TCGA dataset. The average PSI value of tumor samples was subtracted by the average PSI value of normal samples (PSI_Tumor_ – PSI_Normal_) in each cancer type to represent the percent difference in exon 9 or exon 3 PSI value. In total Numb expression, average total Numb transcript level of tumor samples was divided by average total Numb transcript level of normal samples in each tumor type. [(Tumor/Normal) – 1] in percentage was used in histogram to reflect the percent difference in total Numb expression. Wilcoxon test between tumor and normal samples of each cancer type with p-value less than 0.05 is considered significant and marked by an asterisk. Differences greater than 20% are indicated with black asterisk. The boxplot comparison of the PSI value and the total transcript level of Numb between tumor and normal tissue in each cancer type is shown in Supplementary Fig. S1A-C. **(c)** Hazard ratios for progression-free survival with high versus low levels of exon 9 PSI value or total Numb expression. **Left,** within each cancer type, patients are stratified into exon 9 high (top 10^th^ percentile) and exon 9 low (bottom 10^th^ percentile) subgroups based on exon 9 PSI value of *NUMB* in primary tumor. Progression-free interval (in days) was modeled using covariates of exon 9 PSI subgroup and cancer type in Cox proportional hazards model. **Right,** within each cancer type, patients are stratified into high total Numb expression (top 10^th^ percentile) and low total Numb expression (bottom 10^th^ percentile) subgroups based on normalized total Numb transcript level in primary tumor. Progression-free interval was modeled using covariates of total Numb expression subgroup and cancer type in Cox proportional hazards model. Complete tables of hazard ratios and patient stratification are shown in Supplementary Fig. S2A & C and Supplementary Data 1 & 2, respectively.

In contrast, differences in exon 3 inclusion levels were less than 20% and both increased inclusion and skipping were observed (Fig. 1B, Supplementary Fig. S1B-D). Total expression of Numb was lower in 7/21 cancer types by at least 20%, some of which overlap with the 11 cancer types with increased Exon9in, including cholangiocarcinoma (reduced 31% below normal), lung (adenocarcinoma 33%, squamous cell carcinoma 48%), bladder (22%) and liver (20%) (Fig. 1B, Supplementary Fig. S1C-D).

To examine the relationship between the expression of Exon9in Numb isoforms and patient outcome, we extracted the progression-free interval (PFI; time to a new tumor event or death incidence) from the TCGA Pan-Cancer Clinical Data Resource (TCGA-CDR) (29). For each cancer type, patients whose primary tumor has top 10^th^ percentile and bottom 10^th^ percentile of exon 9 inclusion were grouped into exon 9 high and exon 9 low subgroups respectively (Supplementary Data S1). A significantly higher hazard ratio (HR = 1.23, p = 0.012), indicative of decreased progression free survival, was observed in patients with high exon 9 inclusion compared to patients with low exon 9 inclusion, using cox proportional hazards model with covariates exon 9 PSI subgroups and cancer types (Fig. 1C, Supplementary Fig. S2A). In 13/32 cancer types, the survival curves for the high exon 9 inclusion subgroups exhibited a trend towards worse progression free survival (Supplementary Fig. S2B). In contrast, there was no correlation between progression free survival and total Numb transcript level (Fig. 1C, Supplementary Fig. S2C-D, Supplementary Data S2).

### Greater exon 9 inclusion is associated with lower survival in breast cancer

While reductions in total Numb protein expression have previously been implicated in breast cancer progression (11), the expression of *NUMB* splicing isoforms and correlation with progression has not been examined. Analysis of breast cancer patient subgroups showed that all five subtypes of breast cancer exhibit upregulated exon 9 inclusion whereas exon 3 inclusion level was moderately lower in only the luminal subtypes (Fig. 2A, Supplementary Fig. S3A). Stratification of breast cancer patients into exon 9 high and low subgroups within each subtype revealed that high exon 9 inclusion correlates with significantly shorter progression free period (HR = 2.6, p = 0.043) (Fig. 2B, Supplementary Fig. S3B-C) suggesting that the exon 9 PSI value is a potential prognostic marker for breast cancer.

**Figure 2.**
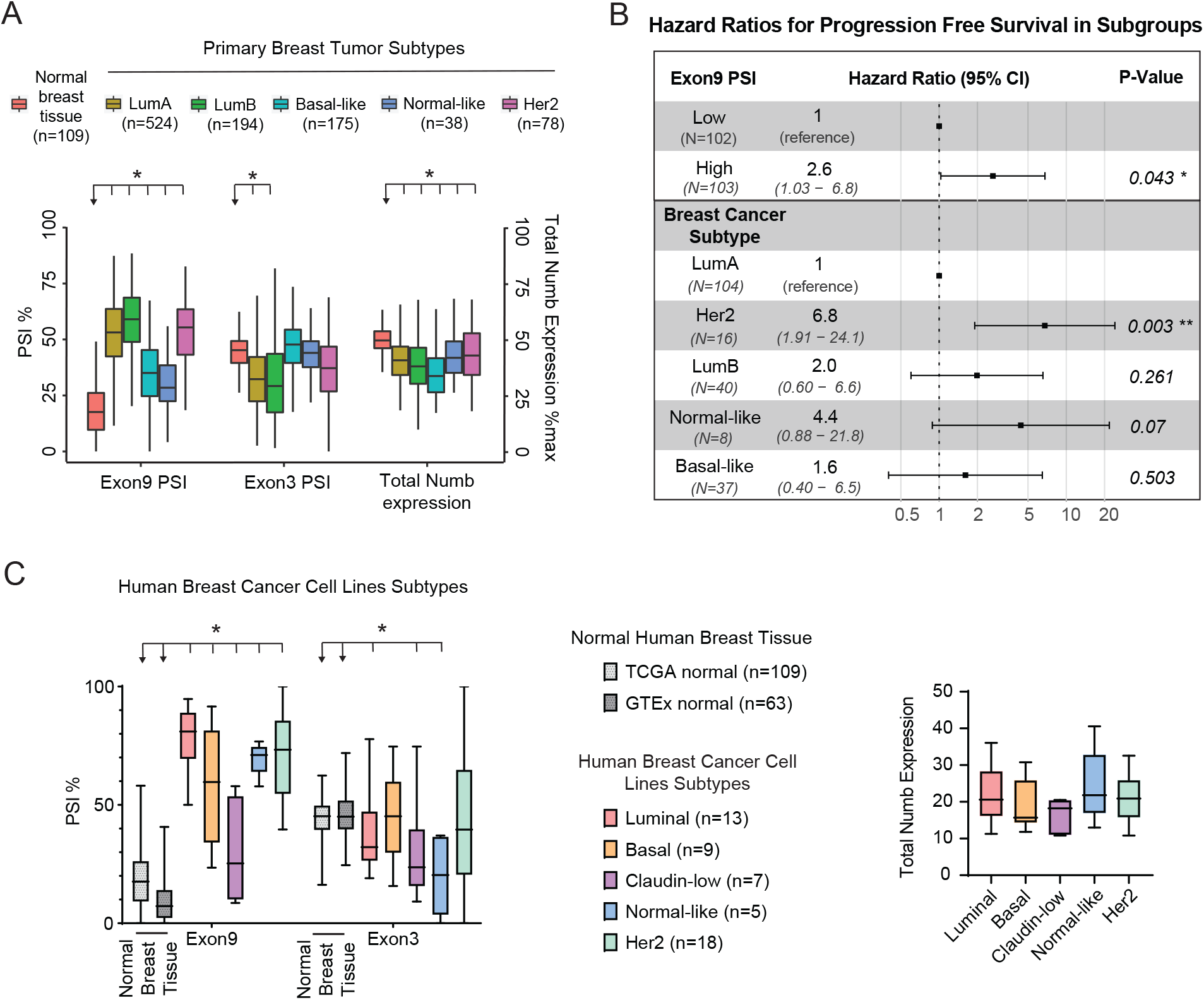
Elevated inclusion of *NUMB* exon 9 in all subtypes of breast cancer is associated with worse prognosis. **(A)** Exon 9 and exon 3 PSI value and total Numb expression in normal human breast tissue and different subtypes of primary breast tumor in the TCGA dataset. Total Numb expression is measured by normalized transcript level in readcounts for Numb scaled by the greatest number. Wilcoxon test was performed between each primary breast tumor subtype and the normal breast samples. *, p-value < 0.05; sample number in brackets. **(B)** Hazard ratios for progression-free survival with covariates exon 9 PSI subgroups and breast cancer subtypes. Within each breast cancer subtype, patients are stratified into exon 9 high (top 10^th^ percentile) and exon 9 low (bottom 10^th^ percentile) subgroups based on exon 9 PSI value of *NUMB* in primary tumor. Progression-free interval (in days) was modeled using covariates of exon 9 PSI subgroup and breast cancer subtype in Cox proportional hazards model. **(C)** Exon 9 and exon 3 PSI value and total Numb expression in different subtypes of human breast cancer cell lines. **Left.** Comparison of exon 9 and exon 3 PSI values for TCGA/GTEx normal breast tissue and human breast cancer cell lines subtypes. Student’s t-test was performed between each breast cancer cell line subtype and TCGA normal or GTEx normal (arrow heads), respectively. *, p<0.05; sample number in brackets. **Right,** Total Numb expression (normalized transcript level expressed as cRPKM) in different subtypes of human breast cancer cell lines.

We also confirmed elevated inclusion of *NUMB* exon 9 using RNA-sequencing data from 52 human breast cancer cell lines (Fig. 2C, Supplementary Data S3) (30) including the basal-like breast cancer cell line MDA-MB-468 (EGFR amplified, PTEN homozygous deletion, p53 mutated) and the claudin-low breast cancer cell line MDA-MB-231 (KRAS, BRAF and p53 mutated) in which prior work has shown increased exon 9 inclusion in response to MEK/ERK pathway activation (23).

### Deletion of Exon9in Numb isoforms impairs breast cancer cell growth

To study the function of Numb Exon9in isoforms in breast cancer, the genomic region of exon 9 was deleted from the MDA-MB-468 cell line using the CRISPR/Cas9 system using three independent sets of guide RNA (gRNA) (Fig. 3A). The three exon 9-deleted (Ex9-del) cell pools generated had diminished expression of the Exon9in Numb isoforms at both RNA and protein level (Fig. 3B) compared to control cell pools (Ctrl) targeted with nonspecific gRNA recognizing a luciferase sequence. Exon 3 inclusion level was not affected by exon 9 deletion (Supplementary Fig. S4A) and targeted pools maintained the mutant population after expansion. Ex9-del clonal lines were also generated, and loss of Exon9in Numb isoforms was confirmed at the protein level (Supplementary Fig. S4B). Targeted clones that failed to delete exon 9 were also used as controls, hereafter referred to as the negatively targeted clone (NTC).

**Figure 3.**
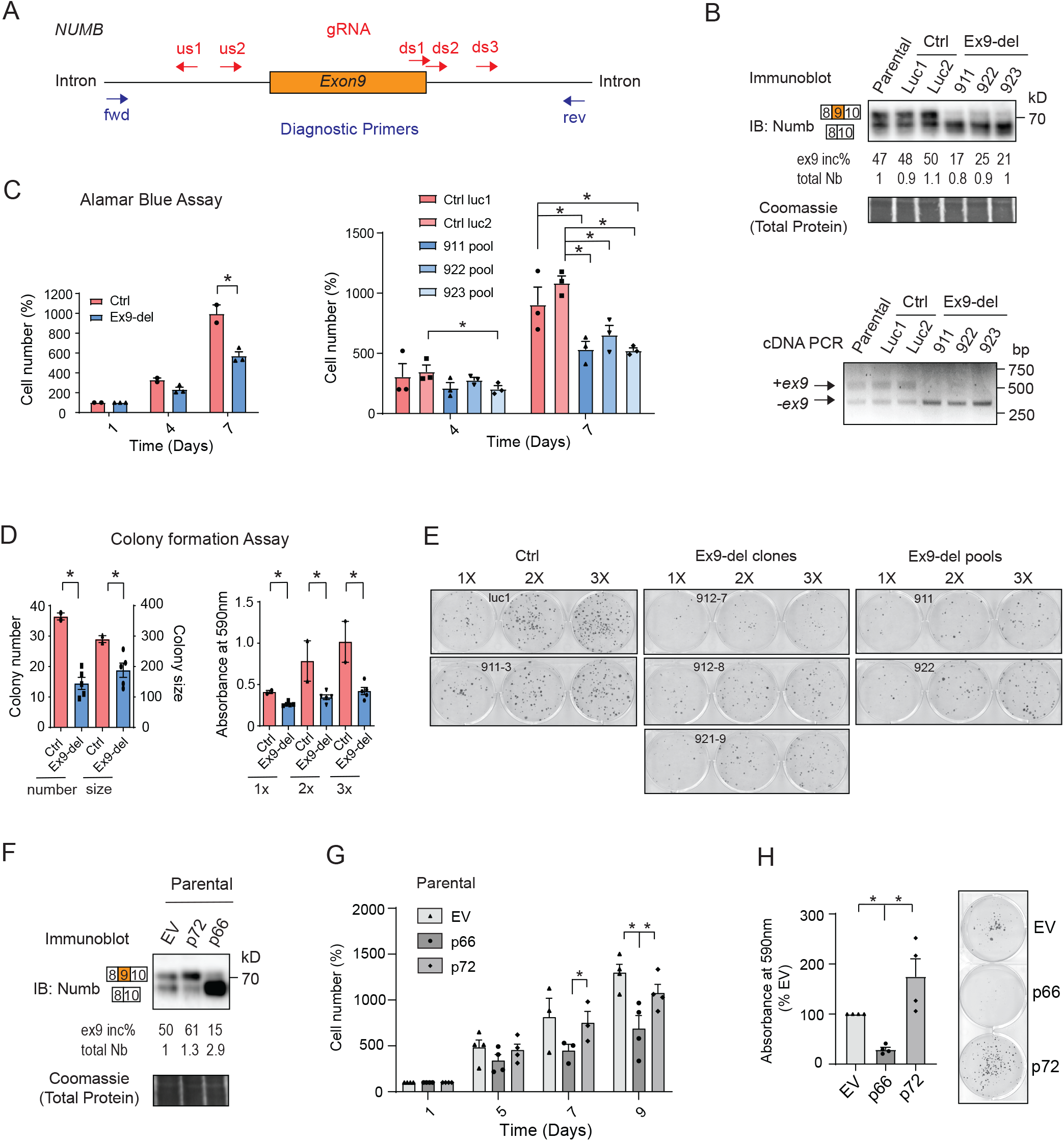
Removal of Exon9in Numb isoforms reduces cell growth. **(A)** Genomic deletion of exon 9 of *NUMB* in human breast cancer cell line MDA-MB-468 cells. Multiple gRNA was designed flanking exon 9 of *NUMB* from the upstream (us) and downstream (ds) intronic regions. Different combination of us and ds gRNA were used to generate 3 different Ex9-deleted (Ex9-del) pools: 911, 922 and 923. Non-specific gRNA against the *Luciferase* (Luc) gene were used as controls (Ctrl). **(B)** The targeting efficiency in Ex9-del pools was confirmed by immunoblot against Numb protein (upper blots) and PCR (lower image) of the exon 9 region from cDNA. The location of the forward (fwd) and reverse (rev) primers used for the diagnostic PCR is illustrated in (A). **(C)** Cell growth rate was measured by Alamar Blue assay for control and Ex9-del cell pools. Cell numbers were normalized to day 1 for each cell pool. Left graph shows Ctrl and Ex9-del pools combined in each group. Each datapoint represents the average of 3 independent experiments. Right graph shows results from each individual pool. Each datapoint represents the average of 3 technical replicates from one experiment. *, p-value < 0.05 by Student’s t-test. Error bar, SEM. **(D)** Colony formation assay. 300 (1×), 600 (2×), and 900 (3×) cells per well for each cell line were seeded and cultured for 3 weeks. Plates were stained with crystal violet to visualize colonies. Colony number and size were quantified by ImageJ (left). Crystal violet was dissolved in 1% SDS solution and the signal for each well was read at 590nm (right). Each datapoint represents the average from 3-4 independent experiments. *, p-value < 0.05 by Student’s t-test. Error bar, SEM. Graphs with individual samples are shown in Supplementary Fig. S4C-D. **(E)** Representative images from single colony formation assay experiment included in data shown in (**D**). **(F)** Stably overexpression of empty vector pcDNA3.1 (EV), p72 or p66 Numb isoform in parental MDA-MB-468 cells. Exon 9 inclusion percentage (e×x9 inc%) is quantified and the total Numb expression was normalized against the first sample. **(G)** Cell growth rate was measured by Alamar Blue assay for overexpression cell pools shown in (F). 4 independent experiments were conducted and each datapoint represents the average of 3 technical replicates from one experiment. *, p-value < 0.05 by pair-wise Student’s t-test. Error bar, SEM. **(H)** Effect of Numb isoform overexpression on colony formation. 900 cells per well for each cell line shown in (F) were seeded and cultured for 3 weeks. Plates were stained with crystal violet and the signal for dissolved crystal violet of each well was read at 590nm. Images from a representative experiment are shown on the right. 4 independent experiments were conducted. *, p-value < 0.05 by Student’s t-test. Error bar, SEM.

To test whether the removal of Exon9in Numb isoforms leads to changes in 2D cell growth, an Alamar Blue assay was performed. The cell doubling rate was decreased by approximately 30% in Ex9-del cells compared to Ctrl cells (Fig. 3C). In colony formation assays, both colony number and colony size were reduced in Ex9-del clones or pools when compared to either the NTC or Luc1 control cells (Fig. 3D-E, Supplementary Fig. S4C-D). We also examined the effect of stable overexpression of the exon 9 containing Numb p72 isoform, and the exon 9 skipped p66 isoform on cell proliferation in parental cells (Fig. 3F). Expression of the p66 Numb isoform in parental cells led to reduced cell doubling rate and colony formation ability compared to cells with the control vector, while p72 Numb overexpression had minimal effect (Fig. 3G-H). The lack of change in cell growth from p72 overexpression is similar to that observed in several studies (22, 24, 29), while others have reported increased cell growth from p72 overexpression (24, 25, 30), suggesting that the effects observed may be dependent on the ratio of expression of the endogenous isoforms in a particular cell context.

### Exon9in Numb isoforms are required for lung metastasis in a breast cancer xenograft model

To determine the contribution of Exon9in Numb isoforms to the tumorigenic activity of basal-like breast cancer, we injected MDA-MB-468 control and Ex9-del cell pools into the mammary fat pad (mfp) of NSG (NOD/SCID/gamma) mice (Fig. 4A). At end point, tumors were excised, and a portion was analyzed by immunoblot and genomic DNA (gDNA) PCR to confirm lack of Exon9in isoform expression in the Ex9-del group tumors (Supplementary Fig. S5A). There was no difference in the primary tumor size or weight between Ctrl and Ex9-del group tumors (Fig. 4B, Supplementary Fig. S5B). In contrast, a significant reduction in spontaneous lung metastases was observed in mice with primary tumors formed by Ex9-del cells compared to mice with tumors formed by Ctrl cells (Fig. 4C-G). Surface metastases were readily observable in the lung tissue from the Ctrl group, but not the Ex9-del group (Fig. 4C).

**Figure 4.**
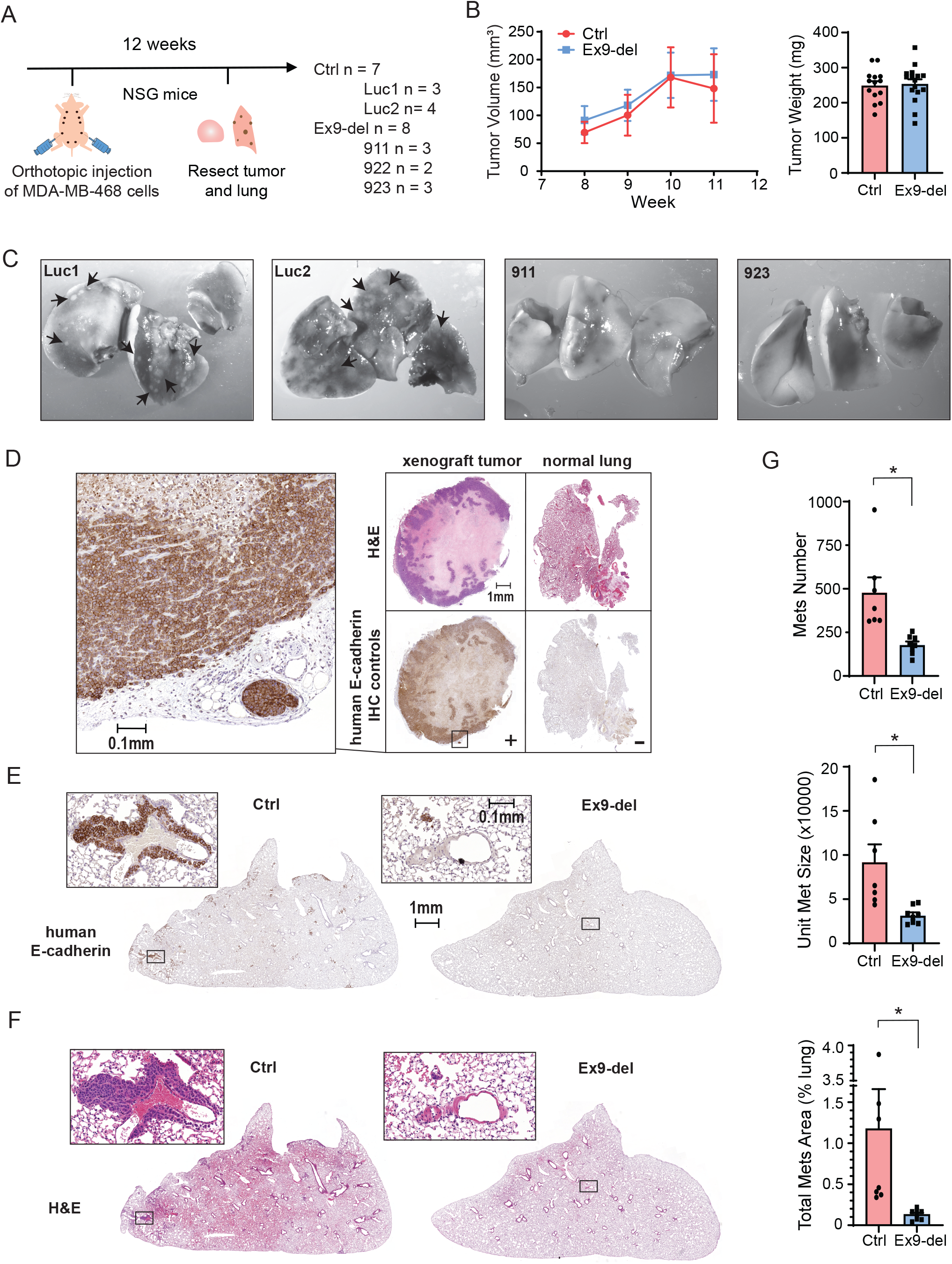
Exon9in Numb isoforms are important for lung metastasis in a breast cancer xenograft model. **(A)** Xenograft experiment design. 3×10^6^ MDA-MB-468 cells were injected into the fourth pair of mammary fat pads (mfp) of NSG mice. For the Ctrl group, a total of 7 mice were injected: 3 were injected with Luc1 cells, 4 were injected with Luc2 cells. For the Ex9-del group, a total of 8 mice were injected: 3 were injected with 911 cells, 2 were injected with 922 cells, and 3 were injected with 923 cells. At endpoint (12 weeks post injection), tumor and lung were resected and analyzed. **(B)** Tumor size and weight measurement. Tumor volume was measured from week8 to week11 by caliper (left). At endpoint the tumor weight showed no difference between Ctrl and Ex9-del groups (right). **(C)** Representative images of surface metastases (arrows) on lungs fixed in Bouin’s solution in Ctrl (Luc1 and Luc2) and Ex9-del (911 and 923) groups. **(D)** Optimization of human E-cadherin antibody. Using MDA-MB-468 xenograft tumor as the positive control (+) and normal lung as the negative control (−), the concentration of human specific E-cadherin antibody was optimized (1:1000) to only pick up signal in positive control but not in negative control (composite image on the right). A magnification of the boxed region in the E-cadherin-stained tumor section was shown on the left. The E-cadherin signal is membrane localized as expected. H&E staining colocalizes actively dividing regions with E-cadherin positive regions in the tumor section. **(E and F)** Representative images of lung sections stained with human specific E-cadherin antibody (E) and H&E (F). E-cadherin staining of metastases appears in brown. In H&E staining metastases are in purple amidst pink background. **(G)** Quantification of lung metastases. Fewer lung metastases, reduced unit metastasis size, and a decrease in total metastases area was observed in lungs from mice injected with Ex9-del cells compared to those injected with Ctrl cells. 2 lobes of lung from each mouse were sectioned at 9 different levels (illustrated in Supplementary Fig. S5D) and stained with the human specific E-cadherin antibody (see Supplementary Methods and Materials section for detailed explanation of quantification steps). The unit metastasis size is calculated by summing the area of individual metastatic sites and dividing that by the number of metastases. Metastases number is normalized to the total area of the lung section. The total metastases area (% lung) is calculated by the sum of metastases area divided by the total lung section area expressed in percentage. Each datapoint represents the average of all sections from one mouse lung. *, p-value < 0.05 by Student’s t-test. Error bar, SEM.

To quantify metastases derived from the primary tumor xenograft, we optimized a human-specific E-cadherin antibody for immunohistochemistry (Fig. 4D). The human E-cadherin positive areas in control xenograft mouse lungs overlapped with actively dividing cells in consecutive sections stained with H&E (Fig. 4E-F), and the E-cadherin signal was localized at the cell membrane (Fig. 4D). While expression of E-cadherin in primary tumor lysate was similar between Ctrl and Ex9-del groups (Supplementary Fig. S5C), areas of E-cadherin staining of lung sections was markedly reduced in the Ex9-del group, indicative of reduced metastasis (Fig. 4E). The total number of metastases per lung, as well as the unit size of metastases and percent area of metastases, was significantly reduced in the Ex9-del group compared to the Ctrl group after sampling at multiple levels (Fig. 4G, Supplementary Fig. S5D).

### Exon9in isoforms regulate EMT and ECM protein networks

To investigate the molecular mechanisms underlying the reduced ability of exon 9-deficient MDA-MB-468 cells to form spontaneous metastases, we compared the total proteomes of two Ctrl and three independently targeted Ex9-del cell pools by quantitative mass spectrometry (MS) using Tandem Mass Tag (TMT) labeled peptides (Fig. 5A). The differential protein abundance was determined by the relative intensities of the TMT tags. Protein abundance across samples showed a strong correlation within Ctrl replicates, while the Ex9-del replicates cluster away from Ctrl (Fig. 5B). Total Numb expression differed by 6.8% between the median signal of Ctrl and Ex9-del replicates, and housekeeping proteins ACTB, TUBA1C and TUBB varied by only 0 – 8% indicating that any differences within 10% were likely due to technical variance (Supplementary Fig. S6A-B, Supplementary Data S4).

**Figure 5.**
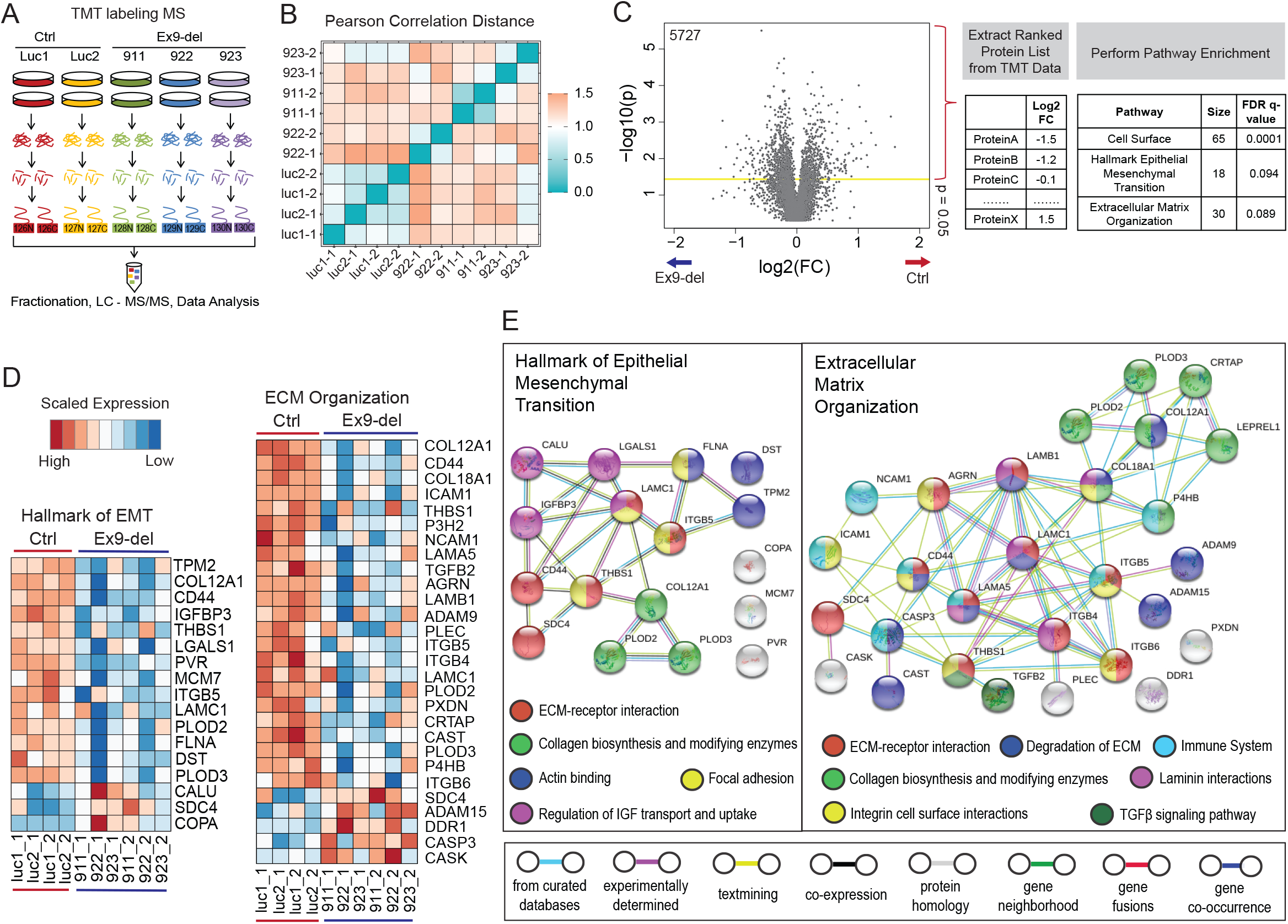
Expression of Exon9in Numb isoforms correlates with elevated level of proteins associated with EMT and ECM Organization. **(A)** Workflow for mass spectrometry (MS) experiment using TMT labelling. Total protein lysate from two MDA-MB-468 Ctrl cell lines and three Ex9-del cell lines were collected. Each line has two technical replicates. **(B)** Pearson Correlation across samples. **(C)** Volcano plot and pathway analysis. Volcano plot was plotted with-log10(p-value) and log2(fold change). P-value was derived from t-test between 4 Ctrl replicates and 6 Ex9-del replicates assuming equal variance with 1 tail alternative. Fold-change (FC) is determined by median of signal for Ctrl replicates divided by median of signal for Ex9-del replicates for 5727 proteins identified. Proteins with p-value < 0.05 were used for pathway enrichment analysis with log2FC as the pre-determined ranking status. Three pathways upregulated in Ctrl samples comparing to Ex9-del samples are highlighted. **(D)** Proteins involved in EMT and ECM organization were upregulated in the presence of Exon9in Numb isoforms. Heatmap showing the scaled (by row) expression level of proteins involved in Hallmark of EMT and ECM Organization pathways which were identified by TMT labeling MS experiment. Proteins with a difference less than 10% between Ctrl and Ex9-del were considered within the noise range and are not shown. **(E)** STRING interaction network of proteins shown in heatmap (**D**). https://string-db.org/

The list of differentially expressed proteins (where Ctrl vs Ex9-del p-value was < 0.05) was subject to pathway enrichment analysis using fold change as the ranking parameter (32). Two metastasis related pathways, Hallmark of EMT and ECM Organization, were identified as significantly altered in the absence of Exon9in isoforms (Fig. 5C, Supplementary Fig. S6C). Proteins involved in these two pathways were generally down regulated in the Ex9-del group compared to Ctrl (Fig. 5D, Supplementary Fig. S6A). Of note, Ex9-del cells had significantly altered expression of membrane proteins with known roles in breast cancer progression and metastasis including CD44 (33), ITGβ4 (34), ITGβ5 (35) and EPHA2 (36) (Fig. 6A-B). Also, the ECM ligand receptor ITGβ5 and the ECM matrix components COL12A1 and LAMC1 were shared between both pathways suggesting that changes in ECM-receptor interactions may be involved in the decreased metastasis seen in Ex9-del cells (Fig. 5E, Supplementary Fig. S6D).

**Figure 6.**
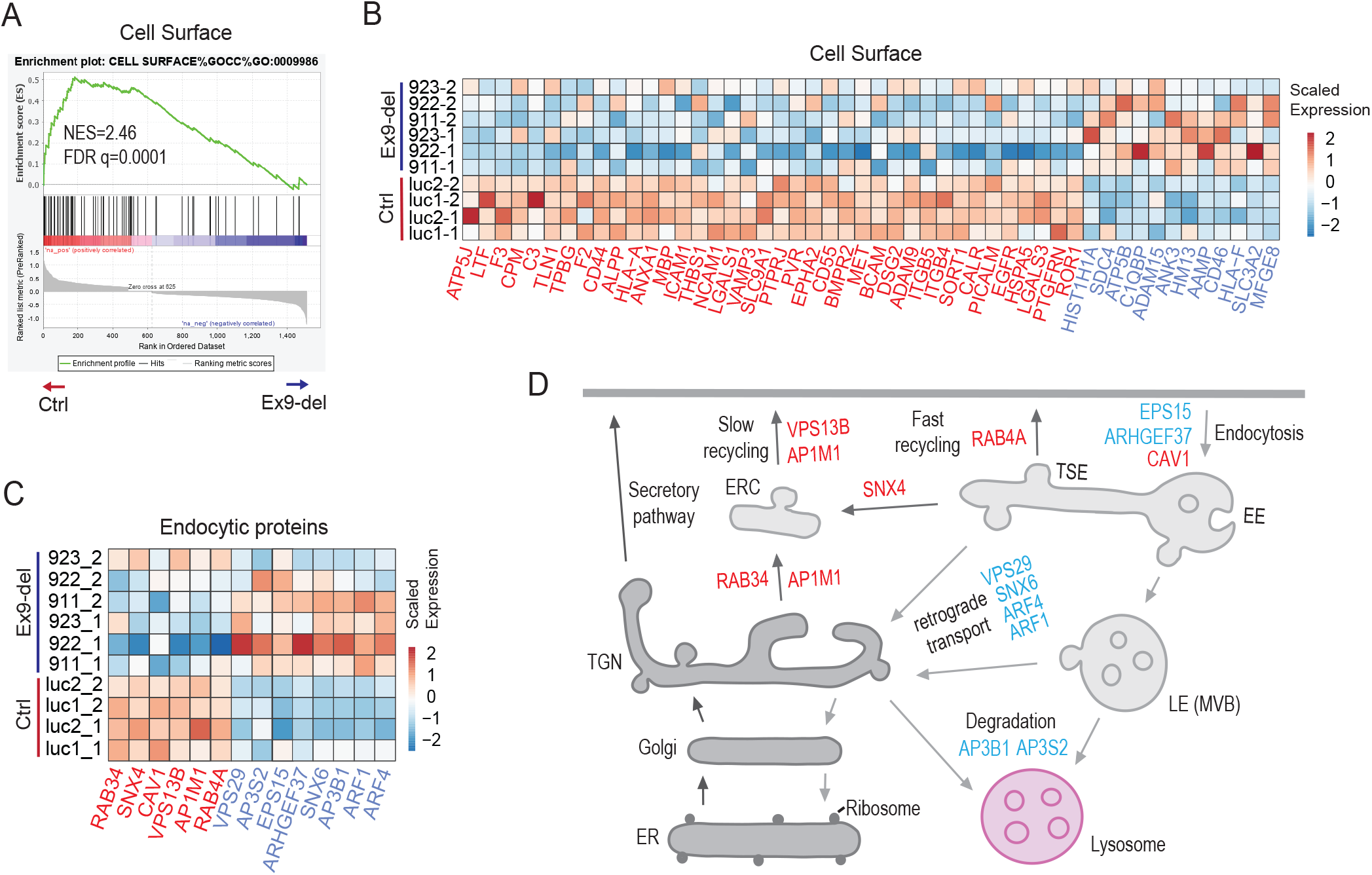
Elevated expression of surface proteins in the presence of Exon9in Numb isoforms. **(A and B)** Enrichment plot (A) and heatmap showing the scaled (by column) expression of proteins involved in the cell surface gene ontology term which were identified by TMT labeling MS experiment of Ctrl and Ex9-del cells. The net enrichment Score (NES) and p-value is shown (A). Proteins with a difference less than 10% between Ctrl and Ex9-del is not shown (B). **(C)** Heatmap showing the scaled (by columns) expression level of proteins involved in endocytosis. Proteins with a difference less than 13% between Ctrl and Ex9-del are not shown. Proteins upregulated in Ctrl samples are in red and proteins upregulated in Ex9-del samples are in blue. **(D)** Illustration of endocytic pathways showing the role of endocytic proteins differentially regulated by Exon9in Numb isoforms. The integration between the endo-lysosomal network and secretory pathway helps to maintain and remodel the cell surface proteome. Following their endocytosis, internalized cell surface proteins enter the early endosome (EE). Selected cargo can be recycled back to the cell surface either directly through tubular sorting endosomes (TSE), termed “fast recycling”, or by transit through the endocytic recycling compartment (ERC), termed “slow recycling”. Recycling back to the cell surface can occur via passage through the trans-Golgi network (TGN) and entry into the secretory pathway, termed “retrograde transport”. Other cargo is selected in the early endosome by the ESCRT machinery to enter the intraluminal vesicles (ILVs) and to be transported on to late endosomes (LE)/multivesicular body (MVB). LE fuses with lysosome to degrade ILVs and their cargo. Proteins identified in **(C)** are placed in their experimentally determined location. Figure adapted from review by Saftig and Klumperman (49)

### Remodeling of endocytic protein network in Ex9-del cells

Numb plays pivotal roles in endocytic trafficking of membrane proteins by clathrin-mediated endocytosis and promoting lysosomal degradation (2, 4). The downregulatory function of Numb is predominantly attributed to isoforms which lack exon 9, while evidence suggests that Exon9in Numb isoforms may antagonize this activity (13). In support of this, the differentially expressed proteins in Ex9-del cells include decreased levels of multiple membrane proteins, suggesting that the absence of Exon9in isoforms may restore Exon9sk mediated lysosomal degradation of cell surface proteins (Fig. 6A-B). Filtering the list of differentially expressed proteins for annotated functions in endocytosis revealed that proteins involved in endocytic recycling and lysosomal trafficking were differentially altered in Ctrl versus Ex9-del cells (Fig. 6C-D). For example, the level of AP3 adaptor subunits, which promote lysosomal trafficking from late endosomes and TGN (37), is elevated in Ex9-del cells, while proteins involved in endocytic recycling and protein secretion (38), such as SNX4, Rab34, Rab4a and AP1 adaptor subunits, are reduced in Ex9-del cells. Furthermore, VPS29, SNX6, ARF1 and AFF4, which all mediate retrograde transportation from endosomes to TGN (38, 39), were increased in the absence of Exon9in Numb isoforms, suggesting that removal of Exon9in might lead to accumulation of surface proteins in intracellular organelles/vesicles in addition to enhanced lysosomal targeting.

### Exon9in isoforms enhance the surface level and signaling of Integrin beta 5

Numb has previously been shown to regulate surface levels of Itgα1 and Itgα5 (6). However, isoform specific functions in relation to integrin regulation have yet to be elucidated. Our proteomic analysis revealed that protein levels of both Itgβ4 and Itgβ5 were reduced in the absence of Exon9in Numb isoforms (Supplementary Fig.S7A). To determine whether Numb isoform expression affected surface levels of Itgβ5 and Itgβ4, Ctrl and Ex9-del cells were incubated with cell-impermeable sulfo-NHS-SS-biotin and biotinylated membrane proteins were isolated using streptavidin-coated Sepharose beads and immunoblotted to determine the relative amounts of Itgβ5, Itgβ4 (Fig. 7A). Whereas loss of Exon9in Numb isoforms had no effect on either surface or total level of the transferrin receptor TfR, the surface level of Itgβ5 was significantly reduced in Ex9-del cells. No change in surface Itgβ4 was observed, although there was a decrease in the total Itgβ4 protein level (Fig. 7B,C). We further tested the level of Itgβ5 and Itgβ4 in Numb isoform overexpressing cell lines. In agreement with the predicted differential isoform effects, total levels of both Itgβ5 and Itgβ4 were higher in cells overexpressing the exon 9 containing isoform, p72, compared to p66 which lacks exon 9. In contrast, overexpression of p66, but not p72 or empty vector, reduced surface levels of Itgβ5 (Supplementary Fig. S7B,C). Numb and Itgβ5 were observed to co-localize in plasma membrane patches in MDA-MB-468 using TIRF microscopy (Fig. 7D). In agreement with surface biotinylation results, both the intensity and surface area of Itgβ5 staining was significantly reduced in Ex9-del cells while Numb plasma membrane localization was unchanged (Fig. 7E, Supplementary Fig. S7D). As well, the proportion of Numb colocalizing with Itgβ5 in plasma membrane patches was reduced in Ex9-del cells (Fig. 7F, Supplementary Fig. S7D). These results are in keeping with a role for Exon9in isoforms in integrin recycling to the plasma membrane while Exon9sk isoforms promote down regulation.

**Figure 7.**
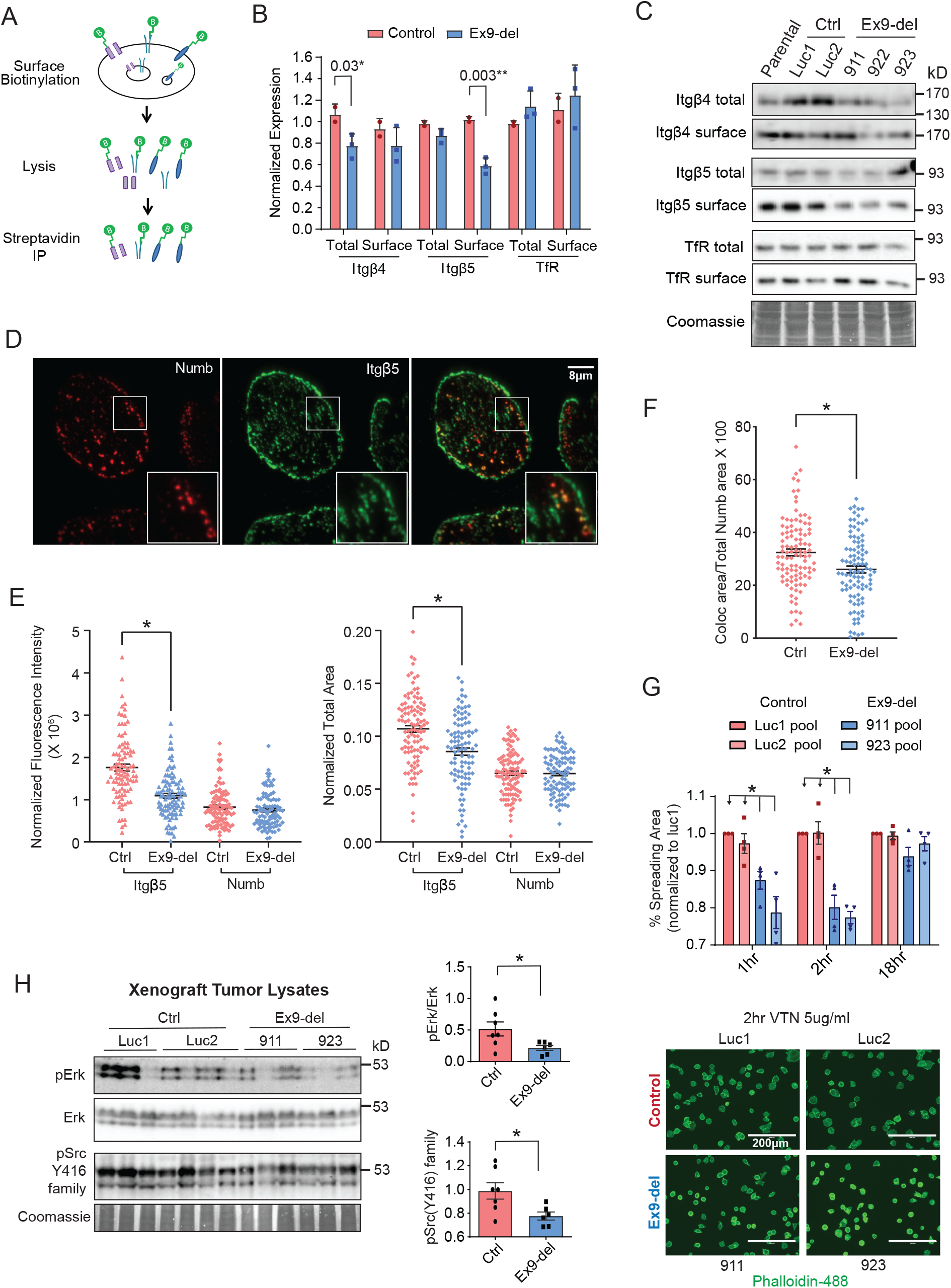
Exon9in maintains surface ITGβ5 and downstream signaling. **(A)** Schematic showing the cell surface biotinylation experiment workflow. Cell surface proteins were labelled with cell-impermeable EZ-Link Sulfo-NHS-Biotin. Biotinylated proteins were captured by Streptavidin-Sepharose beads and analyzed by western blot to compare individual protein level at the surface and in total. **(B)** Quantification of surface-biotinylated ITGβ5, ITGβ4 and TfR as described in (A). The surface level of ITGβ5, but not TfR or ITGβ4, is lower in the absence of Exon9in Numb isoforms. The total protein level was also measured in matched lysate samples removed prior to the streptavidin immunoprecipitation. The total level of ITGβ4 is lower in Ex9-del cells. Each datapoint represents the average of 4 to 6 experiments for individual cell line. *, p-value < 0.05 by Student’s t-test. Error bar, SEM. **(C)** Representative immunoblot images showing biotinylated cell surface ITGβ5, ITGβ4 and TfR and their total level in Ctrl and Ex9-del cell lines. **(D)** MDA-MB-468 cells grown on glass coverslips for 18 hours were fixed, immunostained with anti-NbC (Cy3 anti-GP; red) and anti-ITGβ5 (Alexa488 anti-rabbit; green) and imaged by TIRF microscopy. The enlargement of the boxed region shows that a subpopulation of ITGβ5 is colocalized with Numb. **(E)** Quantification of ITGβ5 and Numb at the basal cell surface. TIRF microscopy images were analyzed using Imaris Analysis Software (see Methods) to quantify the amount of ITGβ5 and Numb localized to basal membrane plaques/structures. For each cell, the total ITGβ5 and Numb fluorescence intensity (left) and surface area (right) were normalized to cell area. Each datapoint represents one cell from 4 independent experiments. *, p-value < 0.05 by Student’s t-test. Error bar, SEM. **(F)** Numb colocalization with ITGβ5 is decreased in Ex9-del cells. For each cell, the percent of total surface Numb colocalized with ITGβ5 at the basal membrane was quantified from TIRF images as described in Methods. Each data point represents a single cell from 4 independent experiments. *, p-value < 0.05 by Student’s t-test. Error bar, SEM. **(G)** Cell spreading on vitronectin is decreased in Ex9-del cells. Suspended cells were seeded onto 5ug/ml vitronectin (VTN) coated plates. At 1hr, 2hr and overnight timepoints, the plates were fixed and stained with phalloidin-488 to visualize the cell periphery for area quantification. 4 images were taken for each condition and at least 10 cells were quantified for each image (n>40). 4 independent experiments were conducted. Each datapoint represents the average of the quantified cell areas in each condition. *, p-value < 0.05 by Student’s t-test. Error bar, SEM. **(H)** E×9-del xenograft tumors exhibit less Erk and Src activation compared to Ctrl tumors. Lysates were prepared from xenograft tumor samples, 7 from the Ctrl group (3 from Luc1 and 4 from Luc2 group) and 6 from the Ex9-del group (3 from 911 and 3 from 923 group). Equal amounts of lysate were immunoblotted for pErk and pSrc (Y416) family proteins. Total Erk and total protein were used to normalize the activated proteins to compare between samples. *, p-value < 0.05 by Student’s t-test. Error bar, SEM.

We further tested whether the decreased surface expression of Itgb5 observed in Ex9-del cells caused functional changes. Firstly, we measured cell spreading on the Itgb5 ECM ligand vitronectin. Compared with controls, Ex9-del cells had a decreased rate of cell spreading over 2 hours of cell attachment to vitronectin (Fig. 7G). Next, we compared the activation status of downstream signaling proteins associated with integrin activation. We used phospho-specific antibodies to measure phosphorylation of ERK and SRC, key proteins in the integrin-FAK signaling axis (41), in lysates from Ctrl and Ex9-del tumors (Fig. 7H). Phosphorylation of both ERK and SRC(Y416) was decreased in Ex9-del primary tumors. This evidence suggests that by regulating the localization and availability of integrins, Exon9in Numb isoforms enhance integrin signaling which contributes to tumor progression and metastasis.

## Discussion

Although the existence of multiple Numb isoforms in mammals was shown more than twenty years ago (14), studies of cancer-related functions of Numb have largely focused on total Numb expression, rather than the role of the individual isoforms in tumor suppression. However, since the discovery that alternative splicing of Numb is a frequent event in lung cancer (13), more recent studies have specifically investigated Numb isoform expression and functions (22, 24, 25). Our analysis of the TCGA and GTEx datasets revealed that increased exon 9 inclusion is a common feature of many cancer types and showed for the first time that *NUMB* exon 9 inclusion is significantly increased in uterine corpus endometrial carcinoma, ovarian serous cystadenocarcinoma, prostate adenocarcinoma, testicular germ cell tumors and breast cancer subgroups. Our work also confirms other published results using distinct datasets which showed that *NUMB* exon 9 inclusion is upregulated in cervical squamous cell carcinoma (20), lung cancer (13), breast cancer (31), hepatocellular carcinoma (22), and urothelial carcinoma of the bladder (21). Several of the cancer types in the TCGA dataset exhibited reduced total Numb mRNA level in addition to the splicing changes in exon 9. Whether these two processes are associated is unclear. Previous studies found no difference in mRNA stability of Exon9in versus Exon9sk Numb isoforms (13, 23) indicating that differential stability is not the cause of the observed reduction in total transcript level of Numb in these cases.

This study also indicates that exon 9 inclusion level, not total Numb expression, is a predictor of patient outcome and suggests that Numb exon 9 expression may have utility as a cancer biomarker. Overexpression of Exon9in Numb isoforms is a common feature across breast cancer subtypes and is a predictor of patient outcome in breast cancer. We conclude from experiments using breast cancer cell lines genetically manipulated to remove Exon9in Numb isoforms that the presence of exon 9 increases cell growth rate, and in a mouse model, Exon9in Numb isoforms are important for the development of spontaneous metastasis, a lethal process in cancer.

Our analysis of proteins that were either up- or down-regulated in cells lacking Exon9in Numb isoforms, suggests that the upregulation of Exon9in isoforms in breast cancer may contribute to cancer progression and metastasis through increased integrin-mediated ECM interactions. Cells that express Exon9in Numb isoforms have greater expression of Itgβ4, surface Itgβ5, and ECM components. Work by others has shown the importance of integrins and ECM in facilitating cancer progression. For example, Itgβ4 expression in breast cancer was demonstrated to be restricted to basal-like subtype, and associated with aggressive behavior (34). Furthermore, Itgβ4 expression in a mouse model of mammary tumorigenesis promoted proliferative signaling, tumor invasion and metastasis (42). Similarly, Itgβ5 promotes tumor growth and angiogenesis in basal-like breast carcinoma cells through Src-FAK and MEK-ERK signaling events (41). In a chick embryo metastasis assay, Itgβ5 mediates epidermal growth factor (EGF) stimulated lung metastasis but not primary tumor growth of human pancreatic carcinoma cells via Src and FAK (35). Finally, lower expression of ECM components may render Ex9-del cells less able to maintain a high level of ECM stiffness and ECM-integrin signaling that is important in tumor metastasis (43).

Quantitative proteomics analysis also revealed changes in the abundance of endocytic proteins in the absence of Exon9in Numb isoforms, supporting different roles for individual Numb isoforms in endocytic trafficking. In particular, lower expression of Rab34, may contribute to reduced surface localization of Itgβ5 in Ex9-del cells, due to its known role in integrin recycling and notably, invasion of breast cancer cells (44). Consistent with our observations, a recent study found that overexpression of the Numb isoform p66, which lacks exon 9, but not the Exon9in Numb isoform p72, decreased the surface localization of ALK in HEK293 cells (26). Our evidence strongly suggests that endolysosomal trafficking of Itgβ5, is similarly influenced by Numb isoform expression.

Intrinsic heterogeneity within breast cancer is reflected in the unique transcriptomic fingerprints and requires differential treatment approaches. Based on transcriptomic profiles, breast cancer has been subcategorized into luminal A (ER+, HER2-), luminal B (ER+, HER2+/−), basal-like (ER-, PR-, HER2-), normal-like and the HER2 enriched subtypes (45). The basal-like subtype has limited treatment options while the other breast cancer subtypes have targeted therapies such as hormone therapies for the luminal subtype and HER2 inhibitors for the HER2 enriched subtype. This has fostered efforts to discover actionable molecular targets to treat patients with basal-like tumors (46). The remodeling of protein expression patterns resulting from the alternative splicing of *NUMB* in breast cancer subtypes may provide opportunities for the development of novel targeted cancer therapies. For example, elucidation of the mechanisms that regulate *NUMB* exon 9 splicing could identify strategies to suppress exon 9 inclusion and restore the down regulatory function of Numb. In addition, strategies to target the elevated surface Itgβ5 or other surface proteins, or the associated downstream signaling in tumor cells with high exon 9 inclusion may have potential to diminish tumor metastases.

## Methods

### TCGA and GTEx RNA-Seq Data Analysis

Alternative splicing analysis was conducted on RNA sequencing (RNAseq) samples from TCGA and GTEx databases (28). The PSI values for *NUMB* exon 9 (chr14:73745988-73746132) and exon 3 (chr14:73783097 - 73783130) were extracted from the supplementary tables. The raw readcounts of all aligned genes were downloaded from the supplementary data portal. TCGA and GTEx samples were categorized into different cancer studies and tissue types using sample metadata supplied on the TCGA and GTEx websites. A two-sided unpaired Wilcoxon test was used to test statistical significance of the difference of the PSI value between tumor and normal samples. The TCGA and GTEx raw gene readcounts datasets were combined and normalized using the DESeq2 package. A Wilcoxon test was applied to measure statistical significance of the difference between overall Numb expression in tumor and normal samples within each cancer type. Analysis of breast cancer cell line RNAseq is described in Supplemental Materials and Methods.

### Survival Analysis

Progression-free intervals (time to a new tumor or death incidence) for each of the TCGA samples were downloaded from TCGA Pan-Cancer Clinical Data Resource (29) and used for survival analysis. The same TCGA barcode structure is used for both *NUMB* exon usage and total expression data and the clinical metadata which allowed for integrated analysis of patient outcome with molecular data. Breast cancer molecular subgroups information was acquired from the TCGAbiolink R package. Within each cancer type or breast cancer subtype, primary tumors are stratified into exon 9 high (top 10^th^ percentile) and exon 9 low (bottom 10^th^ percentile) subgroups based on *NUMB* exon 9 PSI values. NUMB total transcript level (normalized readcount) was also used to stratify patients into Numb high (top 10^th^ percentile) and Numb low (bottom 10^th^ percentile) subgroups. A multivariant Cox proportional hazards model was used to compare the progression-free interval (PFI) between the exon 9 high/low, and Numb high/low subgroups among different cancers or breast cancer subtypes. A univariate Kaplan-Meier test was also done on PFI in exon 9 high/low and Numb high/low subgroups.

### Cell Culture

Human breast cancer cell line MDA-MB-468 cells were cultured in Leibovitz L-15 medium (Gibco #21083-027) supplemented with 10% heat-inactivated FBS and 1xpenicillin/streptomycin. The cell line was authenticated based on STR profiling using services from The Centre for Applied Genomics in SickKids. The cultures were maintained at 37°C in 0% CO_2_ humidified atmosphere and regularly tested for Mycoplasma using SickKids Mycoplasma PCR Cell Line testing service. Cell lines were cultured for no more than 8 culture passages or 2 months. Details of Alamar Blue, Colony Formation and Cell Spreading assays are provided in Supplementary Materials and Methods.

### *NUMB* Exon 9 Targeting

gRNA was designed using tools from crispr.mit.edu. gRNA oligo was cloned into the pSpCas9(BB)-2A-GFP (PX458) plasmid (48). The final vectors with different combination of gRNAs were transfected into MDA-MB-468 breast cancer cell line. GFP positive cells were sorted and collected at the SickKids – UHN flow cytometry facility. The targeted pool of cells was either expanded as a pool or seeded at one cell per well in 96 well plates for clonal expansion. Lysates from both pools and clones were collected and analyzed for the disappearance of specific isoforms. MDA-MB-468 human breast cancer cells were transfected with pcDNA3.1 (empty vector), pcDNA3.1-p72 or pcDNA3.1-p66 human Numb construct. 1000ug/ml G418 was added for 1 week to select for stable cell pools. The stable cells were expanded and maintained in 100ug/ml G418 medium.

### Xenograft Experiments

NOD/SCID/gamma mice were bred in house in The Centre for Phenogenomics (TCP) and studies were conducted as approved by the TCP Animal Care Committee. After washing with PBS, 3×10^6^ MDA-MB-468 human breast cancer cells were resuspended in 30μl PBS followed by injection in the fourth mammary fat pads at both sides of eight to eleven weeks old female NOD/SCID-gamma mice. After a palpable tumor had formed, the greatest longitudinal diameter (length), transverse diameter (width) and height of the tumor was measured with a caliper. Tumor volume was calculated by the modified ellipsoidal formula: tumor volume = 0.5 × (length × width × height). After 12 weeks post injection (when the largest sum of tumor volumes for one mouse reaches 1700mm^3^), tumors were removed, cut into halves at the greatest cross section, and one half was fixed in 10% neutralized buffer formalin (NBF) for 2 days. The other half was cut into small pieces and snap frozen for future lysate/gDNA extraction. Pieces of xenograft tumor samples were homogenized using a tissue grinder followed by either lysis in Laemmli sample buffer or standard Phenol/Chloroform DNA extraction procedures. The lungs were collected, two lobes were fixed in 10% NBF for 2 days followed by paraffin embedding. The other lobes were fixed in Bouin’s solution for 2 days, washed in PBS and imaged. Lung metastases were quantified as described in Supplementary Methods and Materials.

### TMT Labeling Mass Spectrometry Data Analysis

Control and Ex9-del MB-MDA-468 cell protein lysates were prepared, labeled with 11-plex TMT reagents, and analyzed by LC-MS as described in Supplementary Methods and Materials. For TMT abundance normalization, total sum of the abundance values for each channel over all peptides identified within a sample was calculated by Proteome Discoverer analysis software. The channel with the highest total abundance was taken as a reference and all abundances in all other channels were corrected by a constant factor per channel, so that at the end the total abundances are the same for all channels. The output datasheet contains normalized expression signal of each protein (row) in each sample (column). Only proteins with at least 2 peptides observed were considered identified. If any sample has a value of NA, the whole row was eliminated. Pearson correlation distance was calculated across all 10 samples. Fold change (FC) of the expression of each protein between Ctrl and Ex9-del was calculated with the median of 4 Ctrl replicates divided by the median of 6 Ex9-del replicates. Student’s t-test assuming same variance and using one tail alternative was performed between the Ctrl and Ex9-del replicates and a p-value was obtained for each protein. A volcano plot was generated using log10(p-value) vs log2(FC). The list of proteins with p-value < 0.05 was ranked by log2(FC) and used as the input for gene set enrichment analysis (GSEA). Details in referenced paper (32). The Gene sets database used for GSEA PreRanked method is curated by the Dr. Gary Bader’s lab (Human_GO_AllPathways_with_GO_iea_May_01_2020_symbol.gmt). Default settings were used with 1000 permutations. Cell surface and two metastasis related pathways were highlighted with FDR < 25% and were more enriched in Ctrl group compared to the Ex9-del group. Heatmap of expression of genes in the Ctrl and Ex9-del replicates were created for each pathway using heatmap function.

### Surface Protein Biotinylation Assay

70% confluent MB-MDA-468 cells were cooled on ice, washed with cold PBS, and labeled with ice cold 0.2mg/ml EZ-Link Sulfo-NHS-Biotin (Thermo, #21331) in biotinylation buffer (154mM NaCl, 10mM HEPES, 3mM KCl, 1mM MgCl2, 0.1mM CaCl2, 10mM glucose, pH7.6) for 1 hour at 4°C. After two washes with cold PBS, cells were incubated with complete medium with 100mM glycine for 5min on ice to quench unconjugated biotin and washed three times with cold PBS. The cells were then lysed with PLC lysis buffer (50mM HEPES, pH7.5, 150mM NaCl, 10% glycerol, 1.5mM MgCl_2_, 1% Triton X-100, 1mM EGTA, 10mM sodium pyrophosphate, 100mM sodium fluoride containing COMPLETE protease inhibitor tablets (Roche Applied Science). Protein lysates were quantified. 60 microgram of total cell protein lysate was set aside. Equal amounts of protein lysate were mixed with immobilized streptavidin-Sepharose beads (Pierce, #20353) overnight at 4°C to isolate biotinylated proteins and then washed one time in PLC lysis buffer and three times in Nonidet P-40 wash buffer (50mM HEPES, pH7.5, 150mM NaCl, 2mM EGTA, 10% glycerol, 1.5mM CaCl2, 1% Nonidet P-40). The bed volume was removed before resuspending in SDS-Laemmli sample buffer. Recovered biotinylated proteins were analyzed by SDS-PAGE and Western blot using indicated antibodies (4).

### Cell Lysis and Immunoblot

Cells were washed on plates with PBS for 2 times, followed by lysing with 1 X Laemmli Sample buffer (63mM TrisHCl pH6.8, 2% SDS, 10% Glycerol). Cell lysate was collected into microfuge tubes, boiled for 10 min, and mixed for 10min. Lysate were then centrifuged for 10min at maximum speed to remove insoluble materials. BCA assay (Pierce) was used to determine protein concentration. A mixture of 2-Mercaptoethanol (final concentration 5%) and bromophenol blue (final concentration 0.001%) was added to the lysate before loading to SDS-polyacrylamide gels. After gel electrophoresis, and protein transfer, the PVDF-membrane was blocked with 1% fish gelatin (Millipore Sigma, #G7765). The membrane was incubated in primary antibody overnight followed by three 10min TBST washes, 1 hour incubation with HRP conjugated secondary antibody, and at least four 15min TBST washes before development with ECL substrates. BioRad Gel Doc was used to capture the signal after addition of ECL substrates. Membranes were either stripped for reprobing or stained with Coomassie Blue for quantification of total protein expression. BioRad Image Lab software was used to quantify the band signal.

### Quantification of TIRF images

TIRF microscopy images were imported into Imaris 9.3 3D/4D Visualization and Analysis Software (Bitplane) for quantification of fluorescence intensity, colocalization and area of surface structures/plaques. Single, unattached cells were randomly selected for imaging. Each channel (anti-Itgβ5 and anti-Numb) was segmented into a surface based on fluorescence intensity, using thresholding parameters which resolved plasma membrane structures > 150nm diameter. Identical parameters were used for every image within an experiment, where images were acquired using the same acquisition settings. The fluorescence intensity and area of each segmented surface were normalized to the area of the cell. Object based colocalization was used to measure the overlap of Itgβ5 and Numb surfaces using the Surface to Surface Colocalization extension. The colocalized area was normalized to the total area occupied by each surface. Four independent experiments were analyzed (n=29,27,33,15 Luc1 control cells and n=25,25,32,15 Ex9-del cells). The two tailed Student’s t-test was used to test for statistical significance.

### Statistical Analysis

Student’s t test (parametric) and Wilcoxon test (nonparametric) were used to determine statistical significance of differences between control and experimental groups. Survival analysis was performed using the Cox proportional hazards model and Kaplan-Meier survival (log-rank) method. P-value is considered significant when ≤ 0.05. Values are reported as mean ± SEM. All analyses were performed using Graph Pad Prism except for the analysis of RNAseq data which are compared and graphed in R.

## Acknowledgements

The authors thank the following: Michael Reedijk and Qiang Shen for assistance with mouse xenograft experiments and SPARC Molecular Analysis for TMT mass spectrometry, Amanda Luck for mouse colony maintenance and caliper measurements, Nesrin Sabha for help with IHC staining, and Kim Lau and Paul Paroutis (The Imaging Facility, The Hospital for Sick Children) for their assistance with TIRF microscopy and imaging analysis. This work was supported with funds from the Canadian Institutes of Health Research to CJM (FRN 106507).

## Author Contributions

AZ, SED, and CS performed experiments and wrote the manuscript; KO and AB performed experiments. CJM supervised AZ, SED, KO, AB and wrote the manuscript.

