## Supplementary Material for "Isoform-specific functions of Numb in breast cancer progression, metastasis and proteome remodeling"

### Supplementary Materials and Methods

#### Human Breast Cancer Cell Lines RNA-Seq Data Analysis.

Raw RNAseq fastq files of 56 human breast cancer cell lines were downloaded from ebi.ac.uk/ena/browser/view/PRJNA210428 (1). Identification and quantification of the AS events in RNAseq samples were done with Vast-tools (2). "Vast-tools align [paired end file 1] [paired end file 2] --sp Hsa –expr" was called upon for alternative splicing analysis using Vast-tools package (https://github.com/vastgroup/vast-tools). Vast-tools was run using High Performance Computing service at Hospital for Sick Children. Information on *NUMB* gene is extracted using Ensemble ID ENSG00000133961 from the output files. Gene expression levels were determined as reads per thousand mappable positions of target transcript sequence per million of reads. This protocol for estimating gene expression is a corrected version of the widely used RPKM (reads per kilobase of target transcript sequence per million of total reads) and is referred to as "cRPKM" (3).

#### Alamar Blue Assay.

2500 cells were seeded in 96 well plates and grown in normal tissue culture conditions. Fresh medium was added every other day. 110ul of the mixture of 10ul Alamar blue reagent (Invitrogen, #DAI1025) and 100ul of medium was added in each well at indicated time point and incubated for 2 hours. Fluorescence was measured using a Molecular Devices VersaMax 190 Plate Reader. A standard curve was derived by measuring the fluorescence (A562) from a known number of cells. The experimental cell numbers were interpolated from standard curve.

**Colony Formation Assay.**

300, 600 and 900 cells were seeded in 6 well plates and cultured under normal tissue culture conditions. Fresh medium was replaced every week. After 21 days, the plates were fixed and stained with crystal violet (0.1%w/v). Plates were scanned and analyzed by ImageJ. The crystal violet stain was then solubilized using 0.1% SDS, and the signal was measured by absorbance at 590nm.

**Cell Spreading Assay.**

150,000 resuspended cells were seeded in 24-well plates precoated with 5ug/ml vitronectin. After the indicated time of attachment, the cells were washed in PBS and fixed with 2% PFA. The fixed cells were permeabilized in 0.1% Tx100 for 5min, blocked in 3% Donkey Serum for 1 hour at room temperature, and stained with phalloidin-488 for 1 hour at room temperature. The cells were washed in PBS and at least four fields were captured on EVOS FL microscope for each sample. At least 10 cell areas were quantified in each image. In total, at least 40 cell areas were quantified for each condition.

**Immunohistochemistry.**

Fixed lung and xenograft tumor samples were paraffin embedded and sectioned at 5μm thickness along the largest surface area. For tumor sections, they were stained with H&E. For lung sections, they were stained with H&E and human E-cadherin antibody, separately. To analyze metastases, paraffin embedded lung tissue blocks containing one large lobe and one small lobe for each mouse were sectioned at 9 different levels with a distance of 80μm between each level to get a random sampling and representation of how metastases distribute throughout the lung. The slides were immune-stained using human specific E-cadherin antibody to only pick up human cell signal from the metastasized tumor cells within mouse lung background. The concentration of the antibody was optimized using the human breast cancer cell xenograft tumor slide as positive control and the normal lung as negative control. Stained slides were scanned with Zeiss Light Sheet Z1 at 20x at the SickKids Imaging Facility.

**Quantification of Lung Metastases.**

To quantify metastases in each group, the border of each metastatic site with at least 5 positively stained cells was manually delineated using Caseviewer software, and the area and number of metastatic sites per section was calculated. The boundary of the lung section was also delineated to determine the “total area” of the lung section. The “unit metastases size” in each section is calculated by summing the area of individual metastatic sites and dividing that by the number of metastases. The average of the unit metastases size calculated in all sections for each mouse was plotted. Since more metastases are associated with bigger lung area, the “number of metastases” in each section is normalized to the total area of the lung section. The average of normalized metastases number in all sections for each mouse was plotted. The “total metastases area (% lung)” in each section is calculated by the sum of metastases area in that section divided by the area of the lung section. The average of the total metastases area (% lung) in all sections for each mouse was plotted.

**TMT Labeling Mass Spectrometry Experiment.**

Two Ctrl cell pools and three Ex9-del cell pools from independent targeting events were each seeded to two different culture plates to allow cells grown to 70% confluency.

Each plate was washed with cold PBS twice followed by lysis with the lysis buffer (0.5M Tris pH 8.0, 50mM NaCl, 2% SDS, 1% NP-40, 1% Triton X-100, 40mM choloroacetamide, 10 mM TCEP, 5mM EDTA). Lysate were sonicated, then boiled for 10 minutes with shaking. After cooling to room temperature, lysate was centrifuged at maximum speed for 5min. Protein concentration were measured. 200ug of protein lysate for each sample was used. 6 volumes of pre-chilled acetone were added to lysate and left overnight at -20C. Protein samples in acetone were centrifuged and acetone was decanted. 100mM TEAB solution was added to dissolve the protein pellet. 5ug Trypsin/Lys-C mixture was added to each protein sample to digest at 37$^{o}$C overnight. The peptide concentration was measured by the absorbance at 280nm and 50ug was labeled with 11-plex TMT reagents according to the manufacturer's instructions (Thermo, #90110). Lyophilized TMT mixes were fractionated using HPLC to obtain 60 fractions followed by LC-MS analysis (4).

**Immunofluorescent Staining and Microscopy.**

Cells grown on uncoated glass coverslips for 18 hours were fixed with 2%PFA in PBS for 15 minutes, washed once with 100mM glycine/PBS and twice with PBS followed by permeabilization for 10 minutes with 0.1% Triton X-100. Cells were blocked using 3% Normal Donkey Serum (company) and incubated overnight at 4C with primary antibody diluted in blocking buffer: guinea pig anti-NbC (1/250) and rabbit anti-B5 integrin (1/250; D23A5, Cell Signaling Technology cat#3629). Cells were then washed and incubated with fluorescent-tagged secondary antibody for 30 minutes at room temperature: Cy3 Donkey anti-Guinea Pig (cat#706-165-148, Jackson Immunoresearch Laboratories Inc.), Alexa488 Donkey anti-Rabbit (cat#A32790, Invitrogen), Cy3 Donkey anti-rabbit (cat#711-165-152, Jackson Immunoresearch Laboratories Inc.), or Al488 Donkey anti-guinea pig (cat#706-545-148, Jackson Immunoresearch Laboratories Inc.). Imaging was carried out in PBS by TIRF microscopy using a Zeiss ELYRA PS1 with a 100x/1.46 TIRF objective, 488 nm (200mW) and 561 nm (200mW) lasers, and an Andor iXon DU867 detector.

#### Supplementary Figure Legends

**Supplementary Fig. S1. Greater *NUMB* exon 9 inclusion is a common feature of cancer.**

**(A-C)** Boxplot of PSI value of exon 9 (**A**) and exon 3 (**B**) and total expression of Numb (**C**) in tumor and normal samples within each cancer type from TCGA dataset. Purple brackets indicate more than 20% difference of the averages between tumor and normal samples. *, $p<0.05$ by Wilcoxon test of tumor and normal samples in indicated cancer type.

**(D)** Table of % difference of exon 9 and exon 3 inclusion level and total Numb expression between the average of tumor and normal samples. Calculation described in Fig. 1B legend. The number of normal and tumor samples is tabulated. Numbers were in red when $p < 0.05$ by Wilcoxon test of tumor and normal samples. Bigger number was with orange shades and smaller number was in blue shades.

**Supplementary Fig. S2. Greater exon 9 inclusion but not total Numb expression is associated with worse prognosis in cancer.**

**(A and B)** Complete hazards ratios for progression free survival with covariates: exon 9 PSI subgroups and cancer types **(A)** or with covariates: total Numb expression subgroups and cancer types **(B)**. Detailed method is described in Fig. 1C legend. The number of patient samples and the PSI value cutoff for exon 9 high and low subgroups is tabulated in Supplementary Data S1. The number of patient samples and the cutoff value for total transcript level of Numb between Numb expression high and low subgroups is tabulated in Supplementary Data S2.

**(C and D)** Kaplan-Meier progression free survival curve between exon 9 high and low subgroups **(C)** or Numb high and low subgroups **(D)** in each cancer type from TCGA dataset.

**Supplementary Fig. S3. Elevated expression of exon9in Numb isoforms in all subtypes of breast cancer is associated with worse prognosis.**

**(A)** Histogram showing the relative expression of exon 9 or exon 3 Numb isoforms and total Numb in different subtypes of breast tumor and normal breast tissue. The average PSI value from each breast tumor subtype was subtracted by the average PSI value of normal breast samples in each breast tumor subtypes to reflect the percent difference in exon 9 or exon 3 PSI value. For total Numb expression, the average total Numb transcript level of breast tumor samples was divided by the average total Numb transcript level of normal breast samples in each breast tumor subtype. [(Tumor/Normal) – 1] in percentage was used to reflect the percent difference in total Numb expression. Wilcoxon test was used to compare tumor and normal samples for each breast tumor subtype and p-value less than 0.05 is marked as asterisk (*).

**(B)** Stratification of patients into exon 9 high and low subgroups in each breast cancer subtypes for survival analysis. Within each subtype, patients were stratified into exon 9 high (top 10$^{th}$ percentile) and exon 9 low (bottom 10$^{th}$ percentile) subgroups based on their exon 9 PSI value. Number of patient samples and PSI value cutoff is tabulated for the survival analysis in Fig. 2B and Supplementary Fig. S2C.

**(C)** Kaplan-Meier progression free survival curve between exon 9 high and low subgroups in each breast cancer subtypes.

**Supplementary Fig. S4. Removal of exon9in Numb isoforms reduces cell growth.**

**(A)** Confirmation of targeting efficiency in Ex9-del pools. PCR of genomic DNA (gDNA) confirmed exon 9 deleted fragment at genomic level (upper gel). PCR of cDNA flanking exon 3 region showed similar ratio of exon3 inclusion between Ctrl and Ex9-del cell pools (lower gel).

**(B)** Confirmation of Numb expression in clones purified from Ex9-del pools. The negatively targeted clone (NTC) served as one of the controls in colony formation assay (Fig. 3D-E) has undergone the same targeting process but failed to delete exon9.

**(C and D)** Quantification of colony number and colony size **(C)** and absorbance at 590nm **(D)** from individual clone or cell pool in colony formation assay. Refer to Fig. 3D.

**Supplementary Fig. S5. Exon9in deletion reduced lung metastasis from a xenograft mouse model.**

**(A)** Confirmation of exon 9 deletion in xenograft tumor lysate**.** gDNA was collected from xenograft tumor and tissue culture (TC)-grown Ctrl and Ex9-del MDA-MB-468 cells. PCR reaction using diagnostic primers shown in Fig. 3A confirms the presence of exon 9 deleted fragments in the Ex9-del tumors. Tumor lysate was immunoblotted with an Exon9in isoform specific antibody to confirm the loss of Exon9in signal in tumor lysates from the Ex9-del group.

**(B)** Quantification of tumor mass**.** Quantification of % tumor mass from H&E staining of xenograft tumor sections (actively dividing region/total area of the section x 100) showed no statistical difference between Ctrl and Ex9-del groups. *, p-value $< 0.05$ by Student's t test. Error bar, SEM.

**(C)** E-cadherin was expressed at similar level in both Ctrl and Ex9-del tumors. Tumor lysate was immunoblotted with anti-human E-cadherin antibody.

**(D)** Schematic diagram showing level sectioning of lung tissue (red/grey) for detection of metastases (orange). 2 lobes per lung were sectioned at a total of 9 levels for better representation of metastases.

**Supplementary Fig. S6. Expression of Exon9in Numb isoforms correlates with elevated expression of proteins associated with EMT and ECM Organization.**

**(A)** The expression of housekeeping genes in TMT labeling MS experiments is not different in Ex9-del cells compared to Ctrl cells. For all four proteins, $p > 0.05$ after one-tail Student's t test. Error bar, SEM.

**(B)** Expression of Numb family proteins Numb and Numblike (paralog of Numb) in TMT labeling MS experiment. *, $p < 0.05$ by one-tail Student's t test. Error bar, SEM.

**(C)** Enrichment plots of proteins involved in the Hallmark of EMT pathway (left) and the ECM Organization pathway (right) which were identified by TMT labeling MS experiment of Ctrl and Ex9-del MDA-MB-468 cells. Net Enrichment Score (NES) and p-value was shown.

**(D)** Expression of extracellular matrix components in TMT labeling MS experiment. *, p-value < 0.05 by one-tail Student t test. Error bar, SEM.

**Supplementary Fig. S7. Exon9in Numb isoforms expression correlates with elevated expression of proteins associated with EMT and ECM Organization.**

**(A)** Expression of ITGβ5 and ITGβ4 in MS-TMT labeling experiment. *, $p\text{-value} < 0.05$ by one-tail Student's t test. Error bar, SEM.

**(B)** Representative immunoblot image showing surface-biotinylated ITGβ5 and total expression of ITGβ5 and ITGβ4 in parental MDA-MB-468 cells stably overexpressing empty vector (EV), p72 isoform, or p66 isoform.

**(C)** Quantification of surface-biotinylated ITGβ5, total expression of ITGβ4 and ITGβ5 from (B). 4 independent experiments were conducted. *, $p\text{-value} < 0.05$ by one-tail paired Student's t-test. Error bar, SEM.

**(D)** Clustering of Numb with ITGβ5 at the basal cell membrane is decreased in Ex9-del cells. Control and Ex9-del cells were cultured on glass coverslips for 18 hours, fixed, immunostained with anti-NbC (Alexa488 anti-guinea pig; green) and anti-ITGβ5 (Cy3 anti-rabbit; red), and imaged by TIRF microscopy. The enlargement of the boxed region within each image shows that a subpopulation of Itgβ5 and Numb colocalize and cluster within patches at the basal membrane, and the amount of overlap is decreased in Ex9-del cells (lower images) compared to Ctrl cells (upper images). Scale bar 10um.

#### Supplementary Tables

**Table S1 List of gRNA sequence**

| gRNA name | Sequence (**PAM**) | Strand | Score |
|---|---|---|---|
| Exon9-us-1 | GTCAGGTGAATGTACAATAC**TGG** | - | 81 |
| Exon9-us-2 | TCTGCTCCCGTTGTAGCTAA**TGG** | + | 80 |
| Exon9-ds-1 | TGCAGCCACATGTTCGGGTA**AGG** | + | 90 |
| Exon9-ds-2 | AGCCACATGTTCGGGTAAGG**TGG** | + | 89 |
| Exon9-ds-3 | CTAGTTCCCACTCAGCTAAC**TGG** | - | 80 |
| Control-LUC1 | CTTCGAAATGTCCGTTCGGT**CGG** | + | 98 |
| Control-LUC2 | ATGATAAACCGGGCGCGGT**TGG** | + | 97 |

**Table S2 List of primers**

| Primer Name | Sequence | Application |
|---|---|---|
| Exon9-gDNA-Diag-fwd | 5’-cctaagacgcagcagctctg-3’ | PCR on gDNA |
| Exon9-gDNA-Diag-rev | 5’-gtcctgttccacagtcctaatgac-3’ | |
| Exon9-cDNA-sqPCR-fwd | 5’-ccaaggctctttccgagg-3’ | Semi qPCR on cDNA |
| Exon9-cDNA-sqPCR-rev | 5’-catctcccagccagcatc-3’ | |

**Table S3 List of reagents**

| Antibody | Company catalogue number |
|---|---|
| Numb antibody (NbC) | In house rabbit–TNPFSSDAQKAFEIEL–cross react with human & mouse Numb |
| Exon9in specific Numb antibody | In house rabbit–PVAMPVRETNPWAHAPDC–cross react with human & mouse Exon9In Numb isoforms only |
| E-cadherin | Rabbit α human ab40772 |
| ITGβ4 | CST4707, MAB2059, Invitrogen PA5-83325 |
| ITGβ5 | CST3629, HPA001820 |
| Transferrin Receptor | Millipore CBL47 |
| p-Src family | CST2101 |
| p-Erk1/2 | CST9101 |
| Erk1/2 | CST9102 |
| Vitronectin | Gibco A14400 |
| Leibovitz L-15 Medium | Gibco 21083-027 |
| Triethylammonium bicarbonate (TEAB) | ThermoFisher 90114 |
| Acetone | Sigma 650501-1L |
| Tris(2-carboxyethyl)phosphine hydrochloride (TCEP) | Sigma C4706-2G |
| 2-chloroacetamide | Sigma C0267-100G |
| Trypsin/Lys-C Mix Mass Spec Grade | Promega V5073 |

**Supplementary Data S1.** Patient stratification based on exon 9 PSI value.

| Cancer Type | # sample | # top 10% | # bottom 10% | PSI top 10% | PSI bottom 10% |
|---|---|---|---|---|---|
| **ACC** | 54 | 6 | 7 | 27% | 0% |
| **BLCA** | 340 | 35 | 34 | 77% | 29% |
| **BRCA** | 1004 | 103 | 101 | 71% | 26% |
| **CESC** | 260 | 26 | 26 | 77% | 31% |
| **CHOL** | 31 | 4 | 4 | 64% | 17% |
| **COAD** | 268 | 27 | 27 | 81% | 48% |
| **DLBC** | 27 | 3 | 3 | 41% | 14% |
| **ESCA** | 147 | 15 | 15 | 81% | 34% |
| **GBM** | 140 | 14 | 72 | 5% | 0% |
| **HNSC** | 463 | 47 | 47 | 69% | 23% |
| **KICH** | 61 | 7 | 7 | 50% | 14% |
| **KIRC** | 360 | 36 | 36 | 29% | 6% |
| **KIRP** | 251 | 26 | 26 | 55% | 19% |
| **LGG** | 481 | 49 | 359 | 2% | 0% |
| **LIHC** | 274 | 29 | 28 | 63% | 15% |
| **LUAD** | 489 | 50 | 49 | 71% | 21% |
| **LUSC** | 491 | 50 | 50 | 59% | 15% |
| **MESO** | 70 | 7 | 7 | 18% | 2% |
| **OV** | 266 | 27 | 27 | 60% | 19% |
| **PAAD** | 168 | 17 | 17 | 65% | 20% |
| **PCPG** | 147 | 15 | 28 | 9% | 0% |
| **PRAD** | 445 | 45 | 45 | 74% | 36% |
| **READ** | 91 | 11 | 10 | 80% | 48% |
| **SARC** | 212 | 23 | 99 | 7% | 0% |
| **SKCM** | 94 | 10 | 12 | 23% | 0% |
| **STAD** | 302 | 31 | 31 | 78% | 34% |
| **TGCT** | 113 | 12 | 12 | 81% | 26% |
| **THCA** | 454 | 46 | 47 | 45% | 18% |
| **THYM** | 82 | 9 | 9 | 18% | 1% |
| **UCEC** | 129 | 13 | 13 | 78% | 30% |
| **UCS** | 39 | 4 | 4 | 52% | 2% |
| **UVM** | 69 | 7 | 13 | 26% | 0% |

**Supplementary Data S2.** Patient stratification based on total Numb expression.

| Cancer Type | # sample | # top 10% | # bottom 10% | transcript level (normalized readcounts) top 10% | Transcript level (normalized readcounts) bottom 10% |
|---|---|---|---|---|---|
| **ACC** | 54 | 6 | 6 | 8440 | 3531 |
| **BLCA** | 340 | 34 | 34 | 7992 | 3913 |
| **BRCA** | 1004 | 101 | 101 | 8769 | 4200 |
| **CESC** | 260 | 26 | 26 | 9403 | 4946 |
| **CHOL** | 31 | 4 | 4 | 9596 | 4335 |
| **COAD** | 268 | 27 | 27 | 9585 | 4885 |
| **DLBC** | 27 | 3 | 3 | 7007 | 4407 |
| **ESCA** | 147 | 15 | 15 | 13848 | 6023 |
| **GBM** | 140 | 14 | 14 | 5647 | 2934 |
| **HNSC** | 463 | 47 | 47 | 11029 | 4979 |
| **KICH** | 61 | 7 | 7 | 14671 | 8087 |
| **KIRC** | 360 | 36 | 36 | 14564 | 6292 |
| **KIRP** | 251 | 26 | 26 | 14127 | 6313 |
| **LGG** | 481 | 49 | 49 | 5323 | 2705 |
| **LIHC** | 274 | 28 | 28 | 10165 | 4587 |
| **LUAD** | 489 | 49 | 49 | 10873 | 5614 |
| **LUSC** | 491 | 50 | 50 | 8841 | 4094 |
| **MESO** | 70 | 7 | 7 | 8797 | 4050 |
| **OV** | 266 | 27 | 27 | 8541 | 3951 |
| **PAAD** | 168 | 17 | 17 | 11074 | 6219 |
| **PCPG** | 147 | 15 | 15 | 6745 | 4245 |
| **PRAD** | 445 | 45 | 45 | 7047 | 4826 |
| **READ** | 91 | 10 | 10 | 9652 | 5030 |
| **SARC** | 212 | 22 | 22 | 8460 | 3392 |
| **SKCM** | 94 | 10 | 10 | 6993 | 3402 |
| **STAD** | 302 | 31 | 31 | 11600 | 6390 |
| **TGCT** | 113 | 12 | 12 | 6952 | 4055 |
| **THCA** | 454 | 46 | 46 | 7859 | 5105 |
| **THYM** | 82 | 9 | 9 | 7386 | 4311 |
| **UCEC** | 129 | 13 | 13 | 9192 | 4196 |
| **UCS** | 39 | 4 | 4 | 5693 | 2957 |
| **UVM** | 69 | 7 | 7 | 6317 | 4381 |

**Supplementary Data S3.** PSI value for exon 9 and exon 3 of *NUMB* and total Numb expression in human breast cancer cell lines.

| **Cell Line** | **Breast Cancer Subtype** | **PSI *NUMB* Exon9** | **PSI *NUMB* Exon3** | **Numb cRPKM** |
|---|---|---|---|---|
| hcc1143 | Basal | 73.58 | 15.72 | 15.11 |
| hcc1599 | Basal | 65.22 | 20.77 | 15.68 |
| hcc1806 | Basal | 54.1 | 56.65 | 14.93 |
| hcc1937 | Basal | 80.26 | 58.45 | 29.25 |
| hcc3153 | Basal | 48.87 | 32.9 | 22.53 |
| hcc70 | Basal | 84.71 | 50.77 | 30.79 |
| mx1 | Basal | 37.57 | 63.54 | 19.04 |
| sum149pt | Basal | 23.94 | 39.59 | 11.8 |
| sum229pe | Basal | 91.55 | 39.5 | 13.71 |
| bt549 | Claudin-low | 10.14 | 25.6 | 11.28 |
| hcc1395 | Claudin-low | 25.27 | 23.65 | 20.52 |
| hcc38 | Claudin-low | 53.57 | 17.88 | 10.8 |
| hs578t | Claudin-low | 8.57 | 39.61 | 10.96 |
| mdamb157 | Claudin-low | 23.48 | 74.63 | 19.59 |
| mdamb231 | Claudin-low | 38.33 | 9.13 | 18.24 |
| sum1315mo2 | Claudin-low | 57.89 | 15.78 | 20.47 |
| 21mt1 | Her2 | 45.32 | 0 | 19.69 |
| 21mt2 | Her2 | 39.62 | 62.03 | 22.89 |
| 21nt | Her2 | 41.85 | 66.89 | 27.08 |
| 21pt | Her2 | 43.97 | 17.93 | 27.61 |
| au565 | Her2 | 71.17 | 14.08 | 32.55 |
| efm192a | Her2 | 73.2 | 24.68 | 22.13 |
| efm192b | Her2 | 75.64 | 39.59 | 17.24 |
| efm192c | Her2 | 85.42 | 14.08 | 24.4 |
| hcc1419 | Her2 | 88.76 | 21.45 | 14.53 |
| hcc1569 | Her2 | 57.89 | 33.03 | 15.71 |
| hcc1954 | Her2 | 74.19 | 39.53 | 28.91 |
| hcc202 | Her2 | 96 | 27.21 | 15.71 |
| hcc2218 | Her2 | 100 | 100 | 10.8 |
| jimt1 | Her2 | 63.27 | 71.29 | 13.57 |
| mdamb361 | Her2 | 85.84 | 57.65 | 18.23 |
| skbr3 | Her2 | 68.29 | 66.23 | 16.58 |
| sum225cwn | Her2 | 73.44 | 64.25 | 25.56 |
| zr7530 | Her2 | 82.35 | 48.87 | 23.48 |
| 600mpe | Luminal | 77.05 | 30.36 | 29.39 |
| bt474 | Luminal | 50 | 28.72 | 17.46 |

| | | | | |
|---|---|---|---|---|
| bt483 | Luminal | 94.74 | 25.92 | 12.23 |
| cama1 | Luminal | 92.92 | 33.93 | 19.44 |
| hcc1428 | Luminal | 86.52 | 39.61 | 20.54 |
| ly2 | Luminal | 67.86 | 56.63 | 27.31 |
| mcf7 | Luminal | 84.87 | 19.04 | 20.62 |
| mdamb134vi | Luminal | 61.76 | 35.91 | 14.77 |
| mdamb175vii | Luminal | 87.1 | 77.79 | 30.87 |
| sum52pe | Luminal | 74.27 | 26.65 | 20.73 |
| t47d | Luminal | 76.92 | 28.17 | 36.08 |
| t47d_kbluc | Luminal | 70 | 45.43 | 26.33 |
| zr75b | Luminal | 92.45 | 52.13 | 11.28 |
| 184a1 | Normal-like | 70.21 | 20.36 | 21.82 |
| 184b5 | Normal-like | 71.05 | 35.77 | 20.79 |
| mcf10a | Normal-like | 57.8 | 7.35 | 25.1 |
| mcf10f | Normal-like | 76.77 | 37.08 | 40.57 |
| mcf12a | Normal-like | 72.09 | 0 | 12.99 |

**Supplementary Data S4.** Normalized expression of proteins in TMT labeling MS experiment. (Only proteins with $p<0.05$ by one tail Student's t-test were included)

### Supplementary Fig. S1

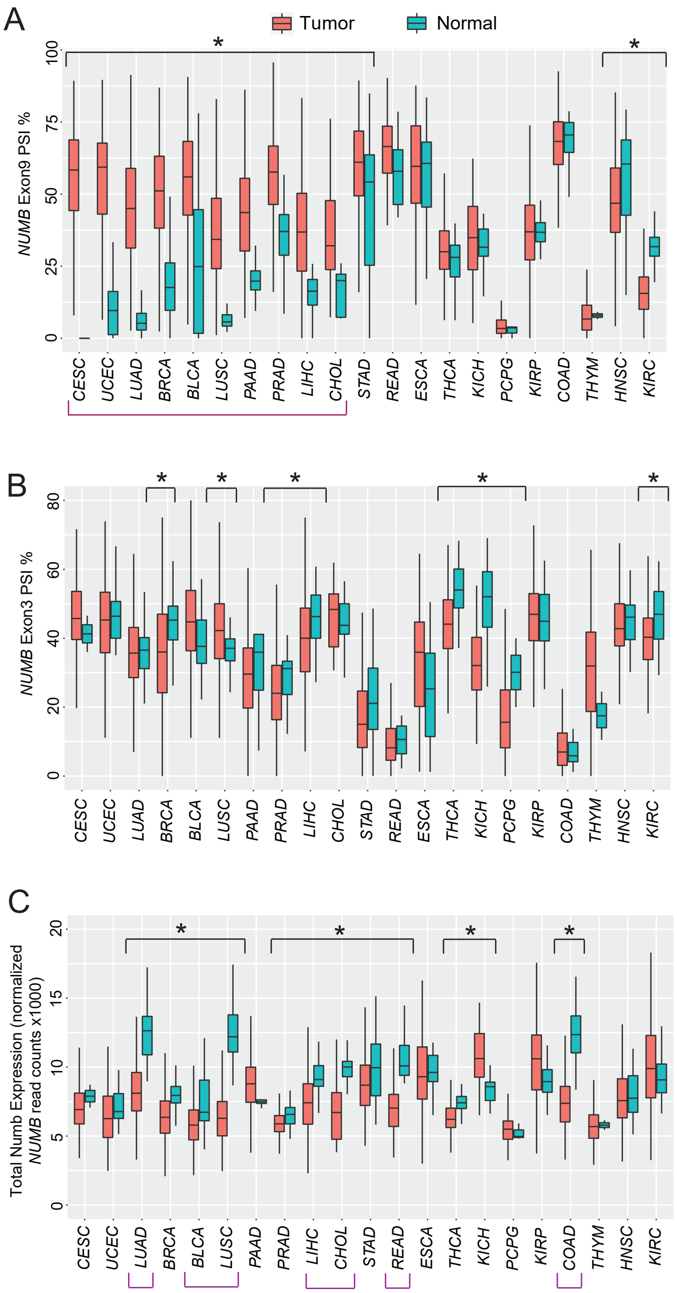


D

| TCGA Study | GTEx normal Tissue Type | # tcga tumor | # tcga normal | # GTEx normal | TCGA tumor vs TCGA normal: ex9 PSI tumor-normal | TCGA tumor vs TCGA normal: ex3 PSI tumor-normal | TCGA tumor vs TCGA normal: Total Nb tumor/ normal-1 | TCGA tumor vs GTEx normal: ex9 PSI tumor-normal | TCGA tumor vs GTEx normal: ex3 PSI tumor-normal |
|---|---|---|---|---|---|---|---|---|---|
| **CESC** | **Cervix** | 260 | 2 | 10 | **0.56** | 0.04 | -0.10 | **0.40** | -0.03 |
| **UCEC** | **Uterine** | 130 | 21 | 32 | **0.44** | -0.03 | -0.08 | **0.54** | -0.03 |
| **UCS** | **Uterine** | 39 | | 32 | | | | **0.20** | -0.03 |
| **LUAD** | **Lung** | 509 | 59 | 122 | **0.39** | 0.01 | **-0.33** | **0.40** | **-0.03** |
| **LUSC** | **Lung** | 497 | 47 | 122 | **0.30** | **0.05** | **-0.48** | **0.31** | **0.03** |
| **OV** | **Ovary** | 267 | | 36 | | | | **0.38** | **-0.17** |
| **BRCA** | **Breast Mammary** | 1010 | 109 | 63 | **0.31** | **-0.09** | **-0.18** | **0.41** | **-0.10** |
| **BLCA** | **Bladder** | 343 | 18 | 11 | **0.28** | 0.06 | **-0.22** | **0.36** | 0.01 |
| **PAAD** | **Pancreas** | 168 | 4 | 65 | **0.23** | -0.02 | 0.17 | **-0.12** | 0.02 |
| **PRAD** | **Prostate** | 446 | 51 | 40 | **0.21** | **-0.04** | -0.09 | **0.32** | **-0.14** |
| **LIHC** | **Liver** | 275 | 46 | 31 | **0.22** | **-0.06** | **-0.20** | **0.11** | **-0.11** |
| **CHOL** | **Liver** | 31 | 9 | 31 | **0.20** | -0.01 | **-0.31** | **0.10** | **-0.06** |
| **STAD** | **Stomach** | 304 | 32 | 75 | **0.14** | -0.04 | **-0.13** | **0.19** | **-0.13** |
| **TGCT** | **Testis** | 125 | | 62 | | | | **0.16** | **0.40** |
| **ACC** | **Adrenal Gland** | 54 | | 52 | | | | **0.08** | **-0.04** |
| **PCPG** | **Adrenal Gland** | 147 | 3 | 52 | 0.02 | -0.12 | 0.05 | **-0.01** | **-0.11** |
| **READ** | **Colon** | 92 | 10 | 79 | 0.07 | 0.00 | **-0.32** | **0.26** | **-0.12** |
| **COAD** | **Colon** | 276 | 40 | 79 | 0.00 | 0.02 | **-0.40** | **0.28** | **-0.13** |
| **ESCA** | **Esophagus** | 147 | 11 | 226 | 0.05 | 0.07 | -0.07 | **0.32** | **-0.12** |
| **SKCM** | **Skin sun exposed** | 94 | | 115 | 0.05 | 0.10 | -0.15 | **-0.15** | **-0.13** |
| **THCA** | **Thyroid** | 454 | 54 | 106 | 0.04 | **-0.10** | **-0.13** | **0.11** | **-0.10** |
| **KICH** | **Kidney cortex** | 62 | 24 | 7 | 0.03 | **-0.18** | **0.39** | -0.03 | -0.14 |
| **KIRC** | **Kidney cortex** | 368 | 72 | 7 | **-0.15** | **-0.07** | **0.11** | -0.21 | -0.07 |
| **KIRP** | **Kidney cortex** | 253 | 32 | 7 | 0.00 | 0.00 | **0.16** | -0.01 | -0.01 |
| **THYM** | | 83 | 2 | | 0.01 | 0.14 | 0.00 | | |
| **GBM** | **Brain** | 143 | | 329 | | | | 0.00 | **0.01** |
| **HNSC** | **Minor Salivary Gland** | 464 | 42 | 5 | **-0.07** | -0.01 | -0.03 | 0.02 | 0.02 |

ACC adrenocortical carcinoma; BLCA bladder urothelial carcinoma; BRCA breast invasive carcinoma; CESC cervical squamous cell carcinoma and endocervical adenocarcinoma; CHOL cholangiocarcinoma; COAD colon adenocarcinoma; DLBC Lymphoid Neoplasm Diffuse Large B-cell Lymphoma; ESCA esophageal carcinoma; GBM glioblastoma multiforme; HNSC head and neck squamous cell carcinoma; KICH kidney chromophobe renal cell carcinoma; KIRC kidney renal clear cell carcinoma; KIRP kidney renal papillary cell carcinoma; LGG brain lower grade glioma; LIHC liver hepatocellular carcinoma; LUAD lung adenocarcinoma; LUSC lung squamous cell carcinoma; MESO mesothelioma; OV ovarian serous cystadenocarcinoma; PAAD pancreatic adenocarcinoma; PCPG pheochromocytoma and paraganglioma; PRAD prostate adenocarcinoma; READ rectum adenocarcinoma; SARC sarcoma; SKCM skin cutaneous melanoma; STAD stomach adenocarcinoma; TGCT testicular germ cell tumors; THCA thyroid carcinoma; THYM thymoma; UCEC uterine corpus endometrial carcinoma; UCS uterine carcinosarcoma; UVM uveal melanoma;

Supplementary Fig. S2

A

**Multivariant Hazard Ratios for progression free survival between exon9 high and low groups**

| Exon9 PSI | | Hazard Ratio (95% CI) | | P-Value |
|---|---|---|---|---|
| Low | (N=1265) | 1 | (reference) | |
| High | (N=804) | 1.23 | (1.047 – 1.46) | 0.012 * |
| **Cancer Type** | | | | |
| ACC | (N=13) | 1 | (reference) | |
| BLCA | (N=69) | 1.33 | (0.550 – 3.21) | 0.528 |
| BRCA | (N=204) | 0.28 | (0.115 – 0.69) | 0.005 ** |
| CESC | (N=52) | 0.93 | (0.357 – 2.42) | 0.88 |
| CHOL | (N=8) | 2.57 | (0.723 – 9.11) | 0.145 |
| COAD | (N=54) | 1.19 | (0.476 – 2.95) | 0.715 |
| DLBC | (N=6) | 0.42 | (0.050 – 3.45) | 0.416 |
| ESCA | (N=30) | 2.21 | (0.837 – 5.81) | 0.11 |
| GBM | (N=86) | 9.28 | (3.994 – 21.55) | <0.001 *** |
| HNSC | (N=94) | 1.12 | (0.468 – 2.67) | 0.802 |
| KICH | (N=14) | 0.27 | (0.055 – 1.35) | 0.11 |
| KIRC | (N=72) | 0.72 | (0.289 – 1.77) | 0.47 |
| KIRP | (N=52) | 0.72 | (0.275 – 1.90) | 0.512 |
| LGG | (N=408) | 1.27 | (0.560 – 2.89) | 0.565 |
| LIHC | (N=57) | 2.08 | (0.860 – 5.03) | 0.104 |
| LUAD | (N=99) | 1.61 | (0.684 – 3.79) | 0.275 |
| LUSC | (N=100) | 0.86 | (0.353 – 2.07) | 0.731 |
| MESO | (N=14) | 3.60 | (1.328 – 9.74) | 0.012 * |
| OV | (N=54) | 2.81 | (1.183 – 6.66) | 0.019 * |
| PAAD | (N=34) | 2.13 | (0.838 – 5.41) | 0.112 |
| PCPG | (N=43) | 0.15 | (0.030 – 0.74) | 0.02 * |
| PRAD | (N=90) | 0.75 | (0.303 – 1.84) | 0.527 |
| READ | (N=21) | 0.69 | (0.211 – 2.27) | 0.544 |
| SARC | (N=122) | 1.55 | (0.667 – 3.60) | 0.308 |
| SKCM | (N=22) | 2.47 | (0.894 – 6.80) | 0.081 |
| STAD | (N=62) | 1.45 | (0.593 – 3.55) | 0.414 |
| TGCT | (N=24) | 0.53 | (0.169 – 1.67) | 0.279 |
| THCA | (N=93) | 0.19 | (0.065 – 0.54) | 0.002 ** |
| THYM | (N=18) | 0.11 | (0.013 – 0.90) | 0.04 * |
| UCEC | (N=26) | 0.69 | (0.231 – 2.05) | 0.502 |
| UCS | (N=8) | 1.77 | (0.500 – 6.29) | 0.375 |
| UVM | (N=20) | 1.94 | (0.688 – 5.47) | 0.21 |

C

**Multivariant Hazard Ratios for progression free survival between Numb expression high and low groups**

| Total Numb Expression | | Hazard ratio (95% CI) | | P-Value |
|---|---|---|---|---|
| Low | (N=797) | 1 | (reference) | |
| High | (N=797) | 0.97 | (0.813 – 1.16) | 0.739 |
| **Cancer Type** | | | | |
| ACC | (N=12) | 1 | (reference) | |
| BLCA | (N=68) | 1.52 | (0.584 – 3.95) | 0.392 |
| BRCA | (N=202) | 0.26 | (0.098 – 0.68) | 0.006 ** |
| CESC | (N=52) | 0.58 | (0.201 – 1.67) | 0.312 |
| CHOL | (N=8) | 2.30 | (0.616 – 8.59) | 0.216 |
| COAD | (N=54) | 0.65 | (0.225 – 1.86) | 0.42 |
| DLBC | (N=6) | 2.81 | (0.544 – 14.52) | 0.217 |
| ESCA | (N=30) | 2.02 | (0.725 – 5.61) | 0.179 |
| GBM | (N=28) | 5.88 | (2.225 – 15.51) | <0.001 *** |
| HNSC | (N=94) | 1.14 | (0.447 – 2.93) | 0.78 |
| KICH | (N=14) | 0.47 | (0.113 – 1.99) | 0.308 |
| KIRC | (N=72) | 0.64 | (0.240 – 1.69) | 0.365 |
| KIRP | (N=52) | 0.93 | (0.345 – 2.51) | 0.888 |
| LGG | (N=98) | 1.36 | (0.534 – 3.45) | 0.522 |
| LIHC | (N=56) | 1.54 | (0.584 – 4.09) | 0.381 |
| LUAD | (N=98) | 1.24 | (0.488 – 3.15) | 0.652 |
| LUSC | (N=100) | 0.82 | (0.315 – 2.11) | 0.673 |
| MESO | (N=14) | 1.90 | (0.622 – 5.83) | 0.26 |
| OV | (N=54) | 2.66 | (1.049 – 6.75) | 0.039 * |
| PAAD | (N=34) | 1.49 | (0.537 – 4.15) | 0.442 |
| PCPG | (N=30) | 0.61 | (0.178 – 2.13) | 0.442 |
| PRAD | (N=90) | 0.46 | (0.168 – 1.24) | 0.123 |
| READ | (N=20) | 0.50 | (0.120 – 2.11) | 0.348 |
| SARC | (N=44) | 1.84 | (0.705 – 4.78) | 0.214 |
| SKCM | (N=20) | 1.32 | (0.402 – 4.34) | 0.646 |
| STAD | (N=62) | 1.22 | (0.456 – 3.24) | 0.696 |
| TGCT | (N=24) | 0.83 | (0.272 – 2.54) | 0.745 |
| THCA | (N=92) | 0.37 | (0.134 – 1.04) | 0.059 |
| THYM | (N=18) | 0.30 | (0.073 – 1.28) | 0.104 |
| UCEC | (N=26) | 0.47 | (0.127 – 1.76) | 0.263 |
| UCS | (N=8) | 1.81 | (0.522 – 6.24) | 0.351 |
| UVM | (N=14) | 1.20 | (0.365 – 3.93) | 0.766 |

B

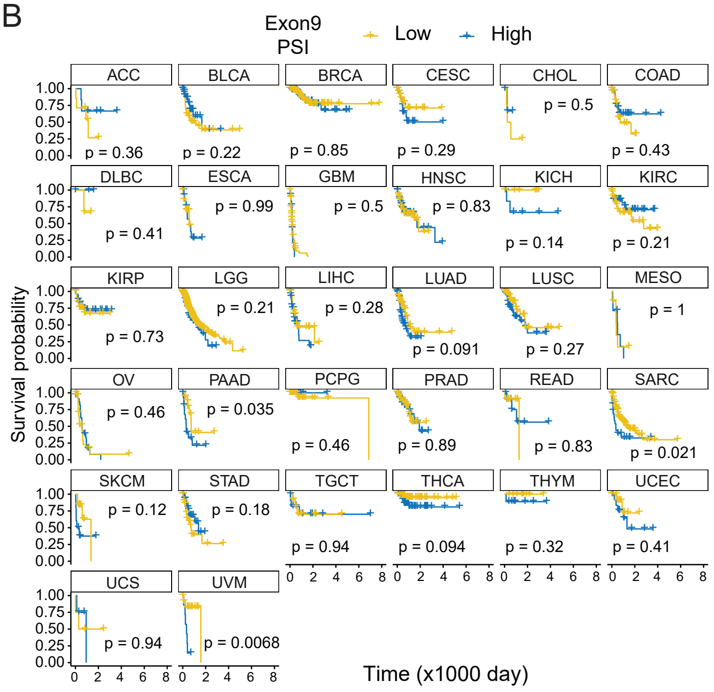


D

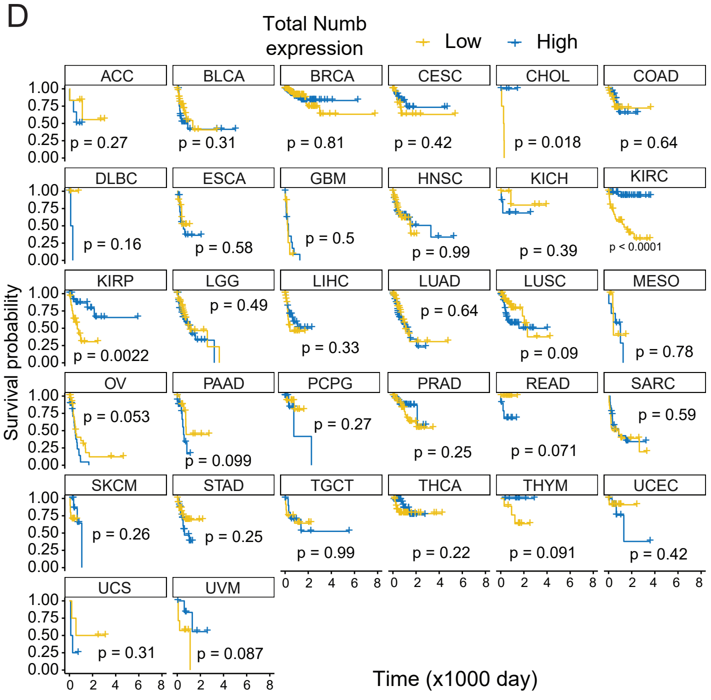

Supplementary Fig. S3

A

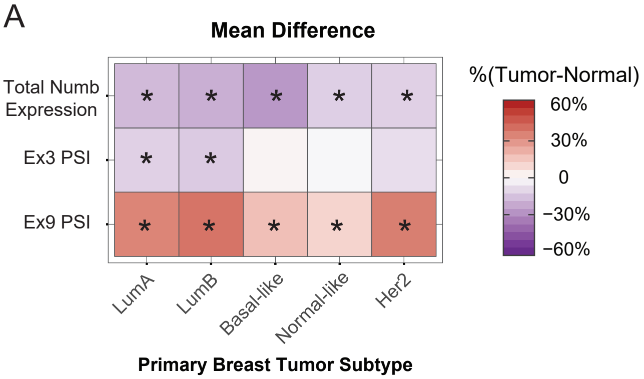


B

**Table 1 Stratification of patients into top 10% exon9 PSI and bottom 10% exon9 PSI group for survival analysis**

| Subtype | # Total Sample | # top 10% | # bot 10% | PSI top 10% | PSI bot 10% |
|---|---|---|---|---|---|
| **LumA** | 516 | 52 | 52 | 72% | 32% |
| **LumB** | 192 | 20 | 20 | 76% | 38% |
| **Basal** | 172 | 19 | 18 | 56% | 15% |
| **Normal** | 38 | 4 | 4 | 43% | 14% |
| **Her2** | 78 | 8 | 8 | 74% | 39% |

C

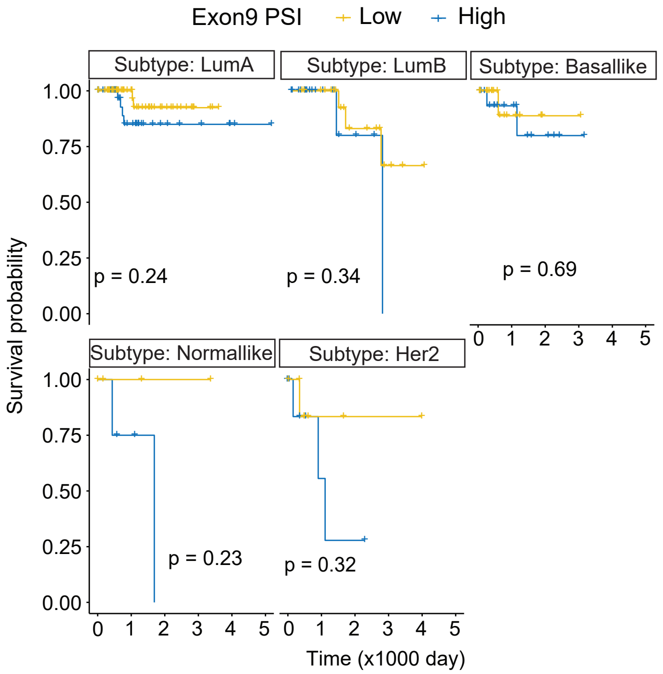

Supplementary Fig. S4

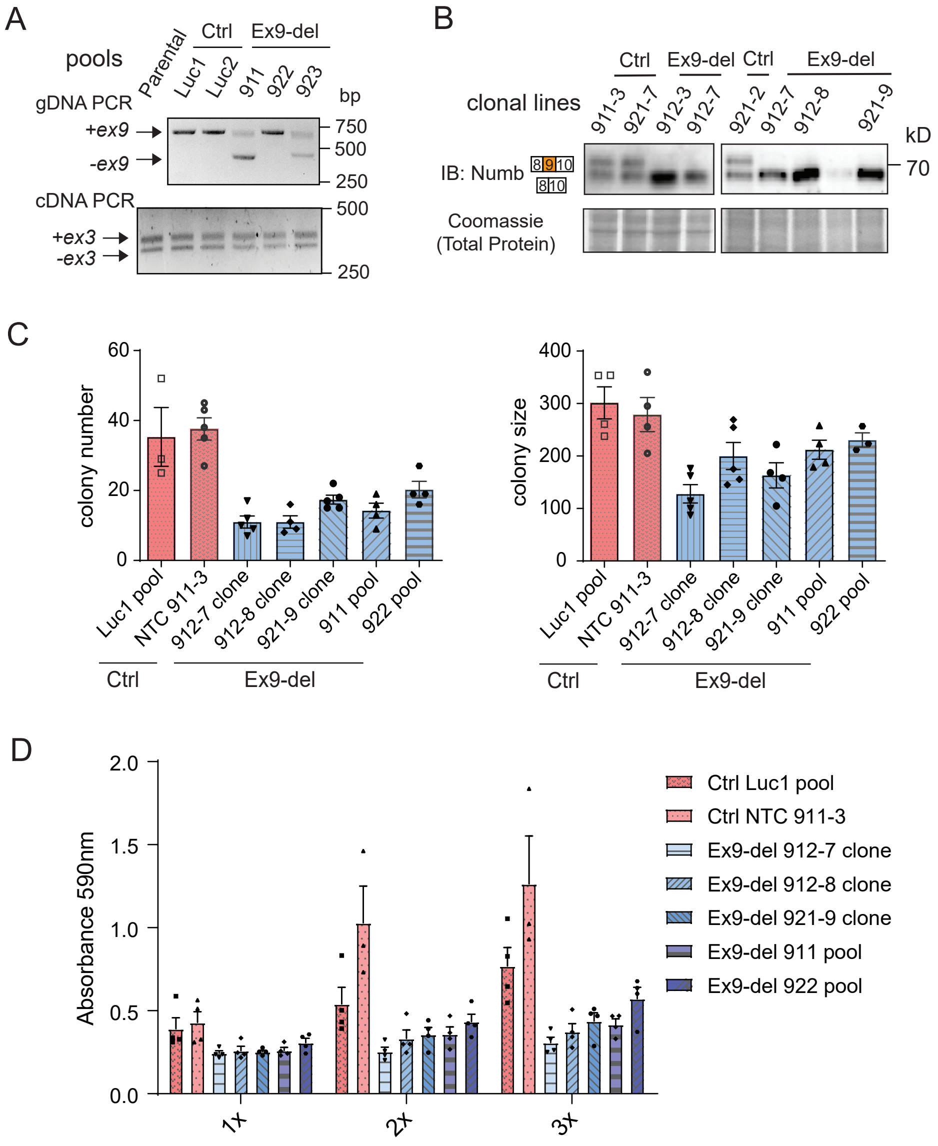

Supplementary Fig. S5

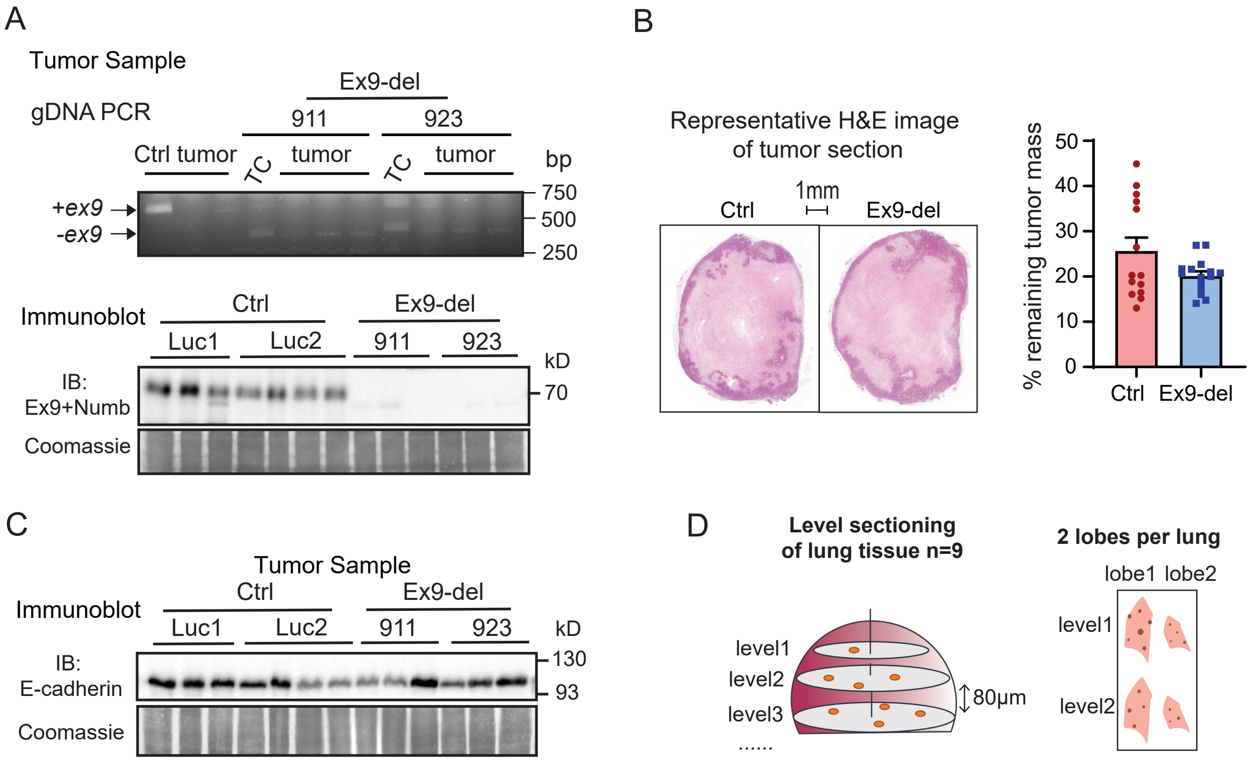

Supplementary Fig. S6

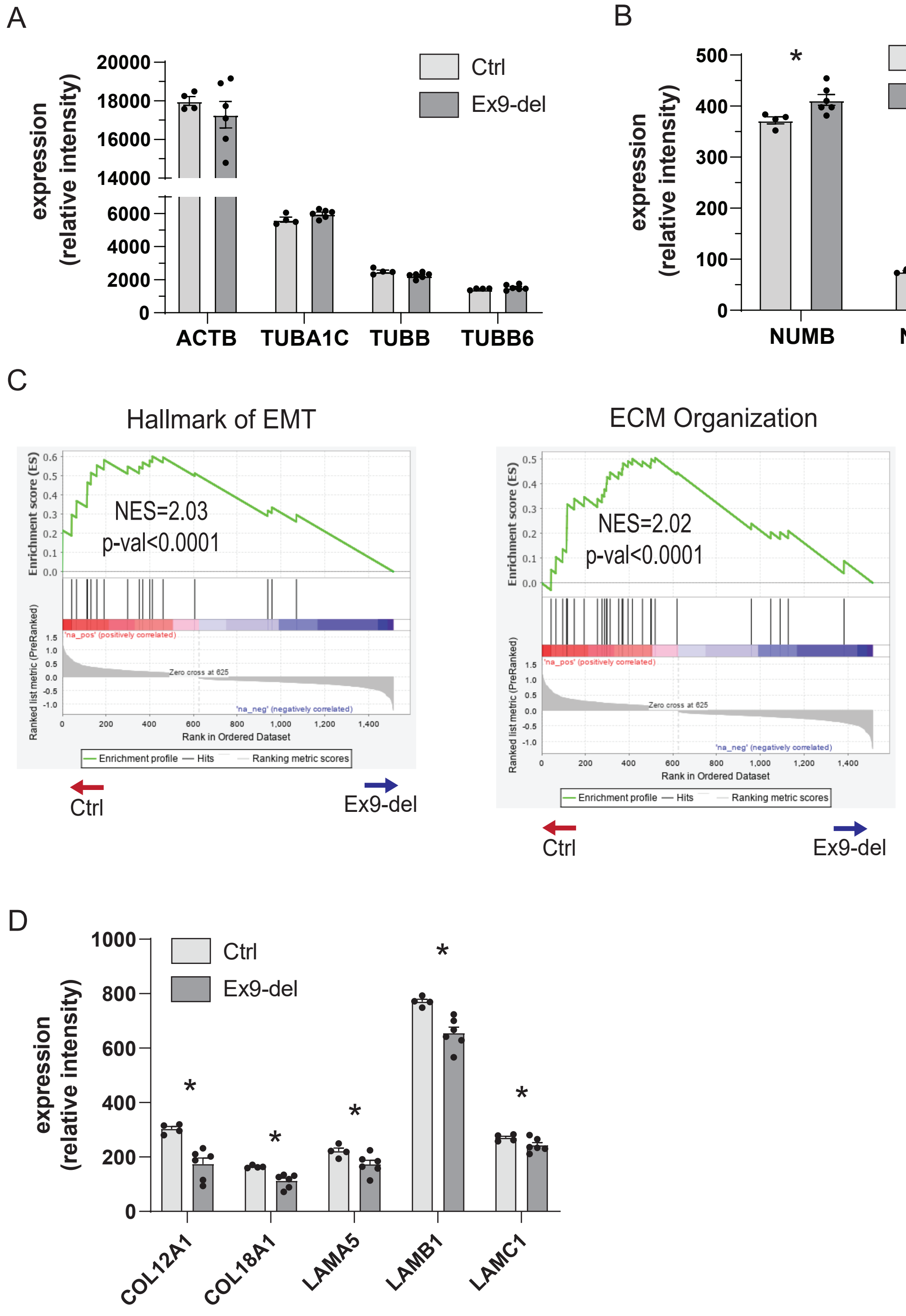

Supplementary Fig. S7

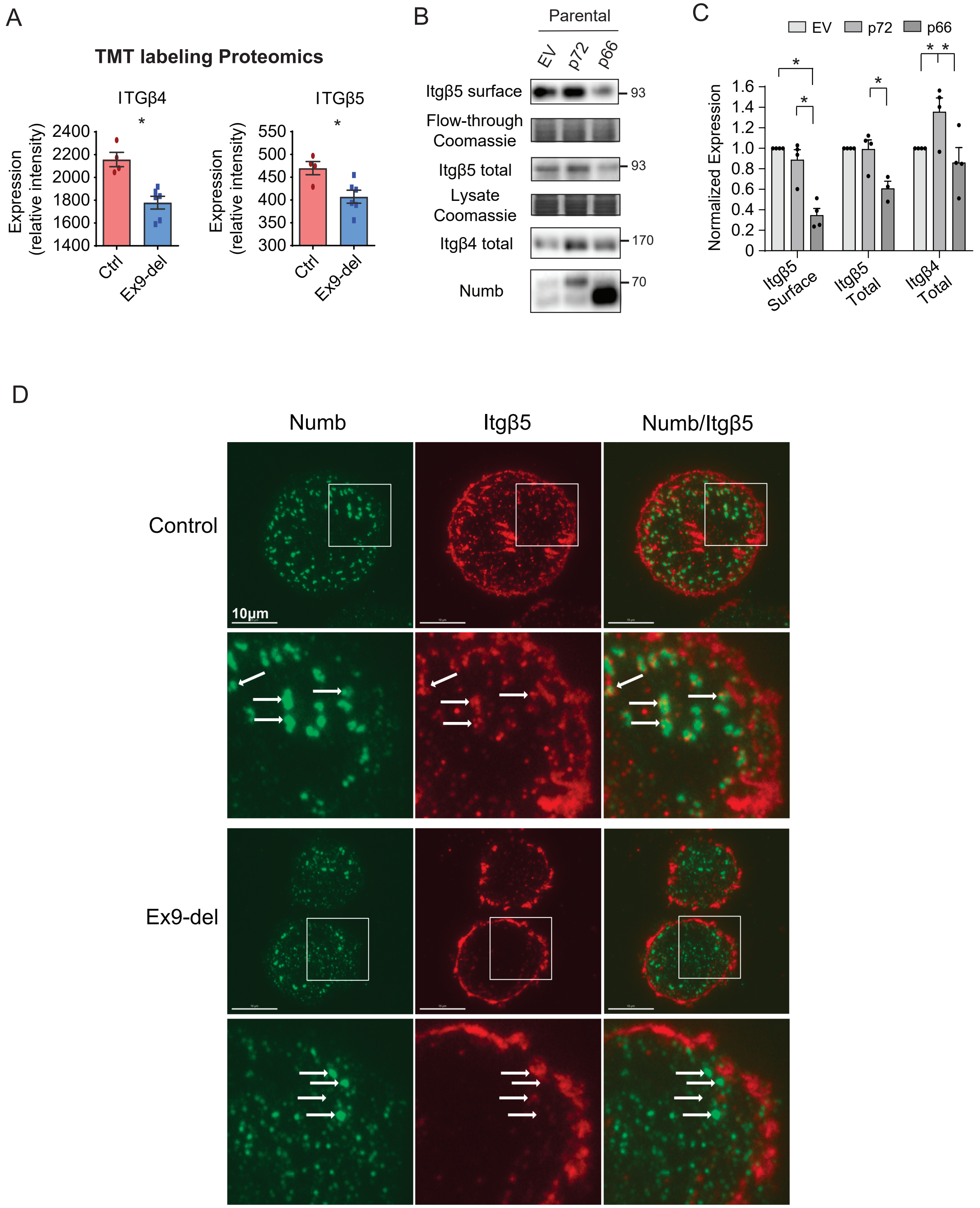
